# Lateral plasma membrane compartmentalization links protein function and turnover

**DOI:** 10.1101/285510

**Authors:** Jon V. Busto, Annegret Elting, Daniel Haase, Felix Spira, Julian Kuhlman, Marco Schäfer-Herte, Roland Wedlich-Söldner

## Abstract

Biological membranes organize their proteins and lipids into nano- and microscale patterns. In the yeast plasma membrane (PM) constituents segregate into a large number of distinct domains. However, if and how this intricate patchwork contributes to biological functions at the PM is still poorly understood. Here, we reveal an elaborate interplay between PM compartmentalization, biochemical function and endocytic turnover. Using the methionine permease Mup1 as model system we demonstrate that this transporter segregates into PM clusters. Clustering requires sphingolipids, the tetraspanner Nce102 and TORC2 signaling. Importantly, we show that during substrate transport, a simple conformational change in Mup1 mediates rapid relocation into a unique disperse network at the PM. Clustered Mup1 is protected from turnover, whereas relocated Mup1 actively recruits the endocytic machinery thereby initiating its own turnover. Our findings suggest that lateral compartmentalization provides an important regulatory link between function and turnover of PM proteins.

## Introduction

The plasma membrane (PM) is an essential barrier, which protects cells from the external environment, orchestrates the exchange of nutrients and metabolites, and serves as a regulatory hub for signaling processes. Lateral segregation of protein and lipid constituents into distinct nano- or microdomains has frequently been observed, and many different models have been proposed to explain the mechanisms for heterogeneous distribution of lipids and proteins within cellular membranes. Prominent examples include the picket-fence model^1^, the lipid-raft hypothesis^2^ and hydrophobic mismatch^3^. However, the extent to which lateral segregation depends on and contributes to specific biological functions at the PM is not clear.

A systematic study of integral membrane proteins in the budding yeast *Saccharomyces cerevisiae* has demonstrated segregation of the yeast PM into a patchwork of many coexisting domains^4^. The observed patterns were proposed to arise through weak protein-lipid interactions^5^. In addition, several distinct PM domains have been previously defined. The stable ‘<u>M</u>embrane <u>C</u>ompartment occupied by (the arginine permease) <u>C</u>an1’ (MCC)^6^ is associated with so-called eisosomes – short furrow-like invaginations that are stabilized by BAR-domain proteins^7^. The MCC and eisosomes have been proposed to play a role in membrane organization, cell-wall morphogenesis and sphingolipid homeostasis^8^. Several tetraspanner proteins such as Sur7 and Nce102, and amino acid transporters including Can1, are recruited to the MCC, where they are thought to be protected from endocytosis^9^. The network-like <u>M</u>embrane <u>C</u>ompartment occupied by <u>P</u>ma1 (MCP), on the other hand, is enriched in the H^+^-ATPase Pma1, which is essential for pH homeostasis and maintenance of the PM potential. Additional domains have been proposed that contain the sterol transporters Lct3/4^10^, the sensor of cell-wall integrity Wsc1^11^ or the Target Of Rapamycin kinase Complex 2 (TORC2)^12^. TORC2 has a wide range of targets, and regulates actin polymerization, endocytosis and sphingolipid synthesis^13^.

Yeast cells that are constantly exposed to fluctuating nutrient levels and environmental stimuli. A large set of cell-surface receptors and nutrient transporters inserted in the PM must be therefore by dynamically modulated. Endocytic downregulation of selected proteins is frequently initiated by ubiquitination of cytosolic lysines via the E3 ubiquitin ligase Rsp5^14^, followed by actin-dependent internalization and degradation within lysosomes^15^. To recognize its targets Rsp5 requires arrestin-related trafficking adaptors, including Art1^16^. Mup1 is a high-affinity, methionine (Met)-specific permease that is ubiquitinated and endocytosed via the Art1/Rsp5 complex in response to substrate excess^17^. More specifically, Met transport by Mup1 has been proposed to induce a conformational change that exposes an N-terminal binding site for Art1^18, 19^. However, while necessary for ubiquitination, this binding site is not sufficient for Art1 recruitment, implying that an additional Art1 interaction site is present in a separate region of Mup1^18^. While ubiquitin-based turnover of plasma membrane proteins is a widespread and well-studied process, it is not clear if and how lateral segregation impacts on endocytic recycling.

In this study we investigated the functional and mechanistic links between PM compartmentalization and membrane protein turnover. Using Mup1 as a model, we show that catalytic activity, ubiquitination and endocytosis of the permease occur within distinct lateral PM domains. In the absence of its substrate Mup1 localizes to distinct clusters at the PM. This clustering depends on a combination of sphingolipids, the tetraspanner Nce102 and TORC2 signaling. Upon Met addition, a conformational switch induces rapid relocalization of Mup1 from the clusters into a disperse network that can be clearly distinguished from other established PM domains. Lateral relocation is accompanied by ubiquitination and endocytic internalization. Importantly, we find that PM clusters provide an environment that protects Mup1 from internalization, while upon relocation, Mup1 actively recruits and re-directs the endocytic machinery.

## Results

### Turnover and PM segregation of the methionine permease Mup1

To investigate a possible connection between PM compartmentalization and protein turnover, we selected the tightly regulated high-affinity methionine permease Mup1 as a model. Upon Met depletion, Mup1 expression was rapidly induced (Figure 1A). The protein then remained upregulated for several hours but increasingly appeared in internal organelles at later time points (Figure 1A). To ensure maximal sensitivity for PM localization experiments, we performed all subsequent experiments after 2.5 h of growth in Met-free medium. At this point Mup1 is one of the most abundant proteins at the PM (Figure 1B). Substrate addition has been shown to induce ubiquitination of Mup1 near its N-terminus, mediated by the arrestin Art1 and the E3 ligase Rsp5^17^. Accordingly, we found that, under our culture conditions, addition of Met induced rapid ubiquitination and degradation of the permease (Figure 1C). Ubiquitination was followed by endocytosis with dose-dependent kinetics. Maximal rates of internalization were observed at Met concentrations exceeding 100 μM (Figure 1D). Deletion of *art1*, truncation of the Mup1 N-terminus (Δ1-52aa = ΔN) or mutation of the ubiquitinated lysine residues^18^ (K27 and K28, designated 2KR from now on)-inhibited Mup1 ubiquitination (Figure 1E) and completely blocked internalization (Figure 1F).

**Figure 1.**
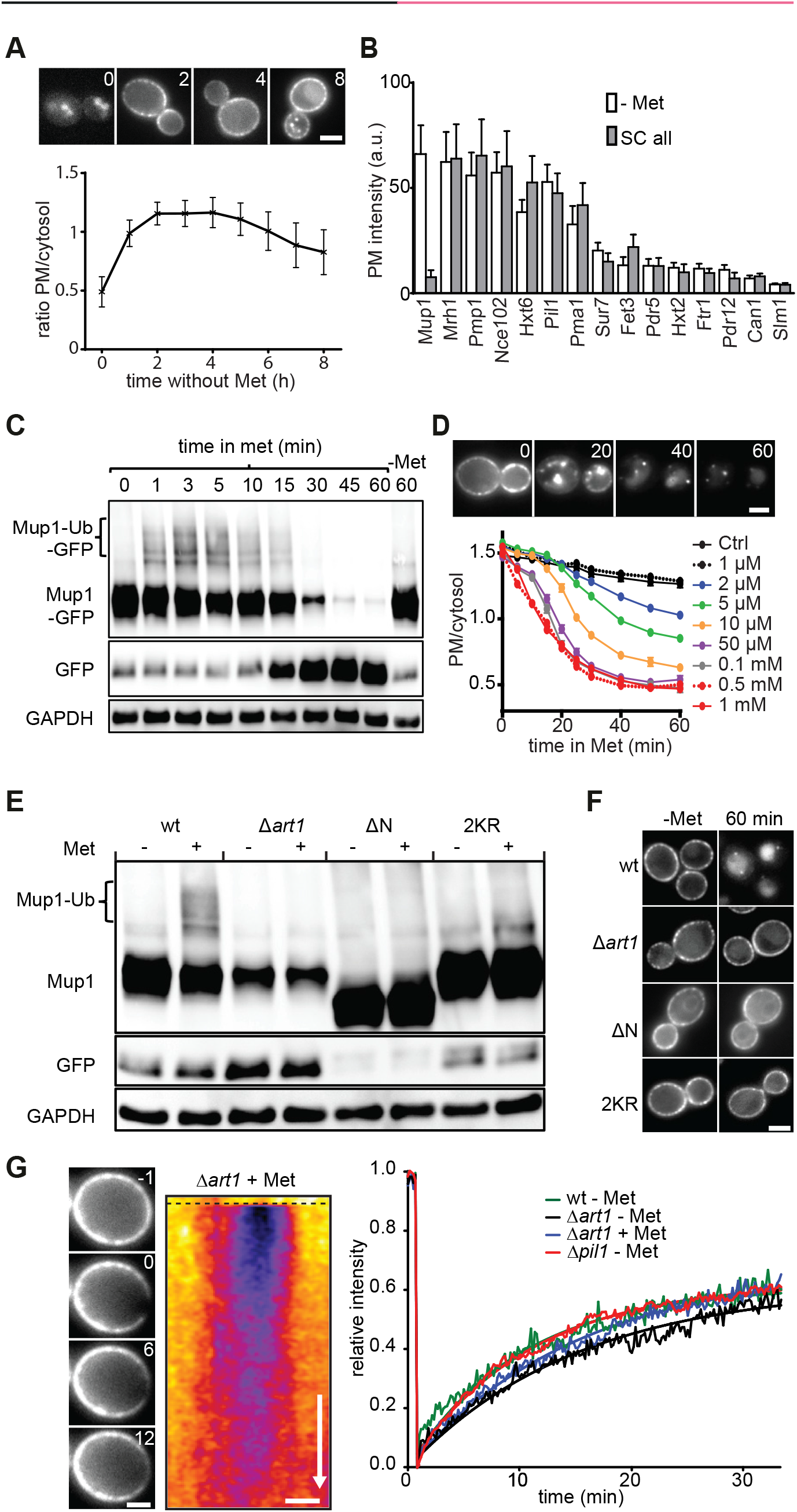
Mup1 is a tightly regulated transporter, which lends itself to the study of the link between PM compartmentalization and protein turnover. (**A**) Expression and localization of Mup1-GFP to the PM upon Met starvation (timestamps in hours). Values are means ± SD, n > 60 cells. (**B**) PM expression of various GFP-fused proteins in cells grown on complete and Met-deficient media. Values are means ± SD, n = 20-300 cells. (**C**) Western blot showing Mup1 ubiquitination (Ub) upon addition of Met. Loading control: glyceraldehyde-3-phosphate dehydrogenase (GAPDH). (**D**) Influence of different Met concentrations on Mup1 internalization. Values are means ± SEM, n > 50 cells (timestamps in min). (**E**) Western blot showing Mup1 ubiquitination (Ub) upon Met addition of the indicated mutants. Ubiquitination requires the α-arrestin Art1 and occurs at lysines K27 and K28 in the cytosolic N-terminal segment of the protein. Samples were prepared 5 min after addition of Met. Loading control as in (C). (**F**) Mup1 endocytosis of the mutants used in (E). (**G**) FRAP analysis of Mup1-GFP (timestamps in min relative to localized bleaching of fluorescence). The kymograph was drawn around the cell periphery and shows fluorescence recovery in the bleached area. Recovery curves represent averaged values from n = 3-5 cells. Scale bars: 2 μm. All values are listed in Table S1.

We had previously reported that integral PM proteins in yeast exhibit rates of lateral mobility that are several orders of magnitude slower than those found in mammalian cells^4^ and this was recently confirmed by single molecule tracking experiments^20^. To determine whether Mup1 also followed this pattern, we performed fluorescence recovery after photo bleaching (FRAP) experiments either in the absence or presence of Met. In agreement with our earlier observations, fluorescent Mup1 diffused into the bleached area of the PM only very slowly, with a *t*1/2 of 5 to 10 min (Figure 1G). Mobility was not affected by ubiquitination, as the kinetics of FRAP recovery were not altered by deletion of *art1* (Figure 1G). Importantly, slow recovery of Mup1 at the PM was partially dependent on the actin cytoskeleton (Fig. S1A), indicating a significant contribution of membrane turnover. The combination of rapid but tightly controlled induction and endocytic removal, together with its low lateral mobility, makes Mup1 an ideal test protein with which to study the link between lateral PM compartmentalization and membrane turnover.

We next analyzed the lateral organization of Mup1-GFP within the PM. Equatorial images already revealed that, in the absence of Met, much of the Mup1-GFP signal at the PM was restricted to defined clusters that strongly colocalized with eisosomes marked by Pil1-RFP (Figure 2A). Profiles across the PM indicated that Mup1 was localized further away from the cell center than Pil1. This localization is even better seen by super resolution microscopy^20^ and is typical for proteins of the MCC, which likely do not enter the furrows formed by eisosomes^8^. By combining total internal reflection fluorescence microscopy (TIRFM) and 2D deconvolution we confirmed the localization of Mup1 to the MCC, but also found weaker staining in a network-like distribution connecting the patches (see asterisk in the lower plot of Figure 2A). The network factor, which is equivalent to the variance of the intensity distribution in our deconvolved TIRFM images, confirms the mixed localization pattern for Mup1. Markers that strongly cluster in PM patches such as Sur7, exhibit a low network factor, whereas proteins distributed in network-like patterns such as Pma1 show higher values. For Mup1 an intermediate value was obtained (Figure 2B). As expected for an MCC marker, Mup1 completely lost its clustered distribution upon deletion of *pil1* (Figure 2B), while its lateral mobility was essentially unchanged (Figure 1G). Interestingly, when Met was added to the medium, Mup1 moved out of the MCC patches and its pattern became more diffuse (Figure 2C, D). However, due to strong signal from endosomes labeled with internalized Mup1, it was difficult to accurately follow the pattern of Mup1 localization for more than 15 min after Met addition (Figure 2D). We therefore performed more detailed quantitative analyses using the internalization mutants mentioned above (*Δart1* background, ΔN and 2KR, Figure 1F). In all three mutants, Mup1-GFP exhibited a clustered distribution under Met starvation, but became much more uniformly distributed after substrate addition (Figure 2E). Using an automated algorithm to analyze colocalization in thresholded TIRFM images (Figure S1B), we confirmed Mup1 exit from the MCC upon substrate addition, illustrated by the decreasing colocalization with Pil1 (Figure 2F). Importantly, in all three internalization mutants, this relocation was completely reversible upon washout of Met (Figure 2G). Our results thus show that, in the absence of its substrate, Mup1 is clustered in the MCC domain. This localization is independent of ubiquitination (*Δart1*, 2KR) or other N-terminal signals (ΔN). Upon addition of Met, Mup1 rapidly relocates into a network-like domain, becomes ubiquitinated and is subsequently internalized.

**Figure 2.**
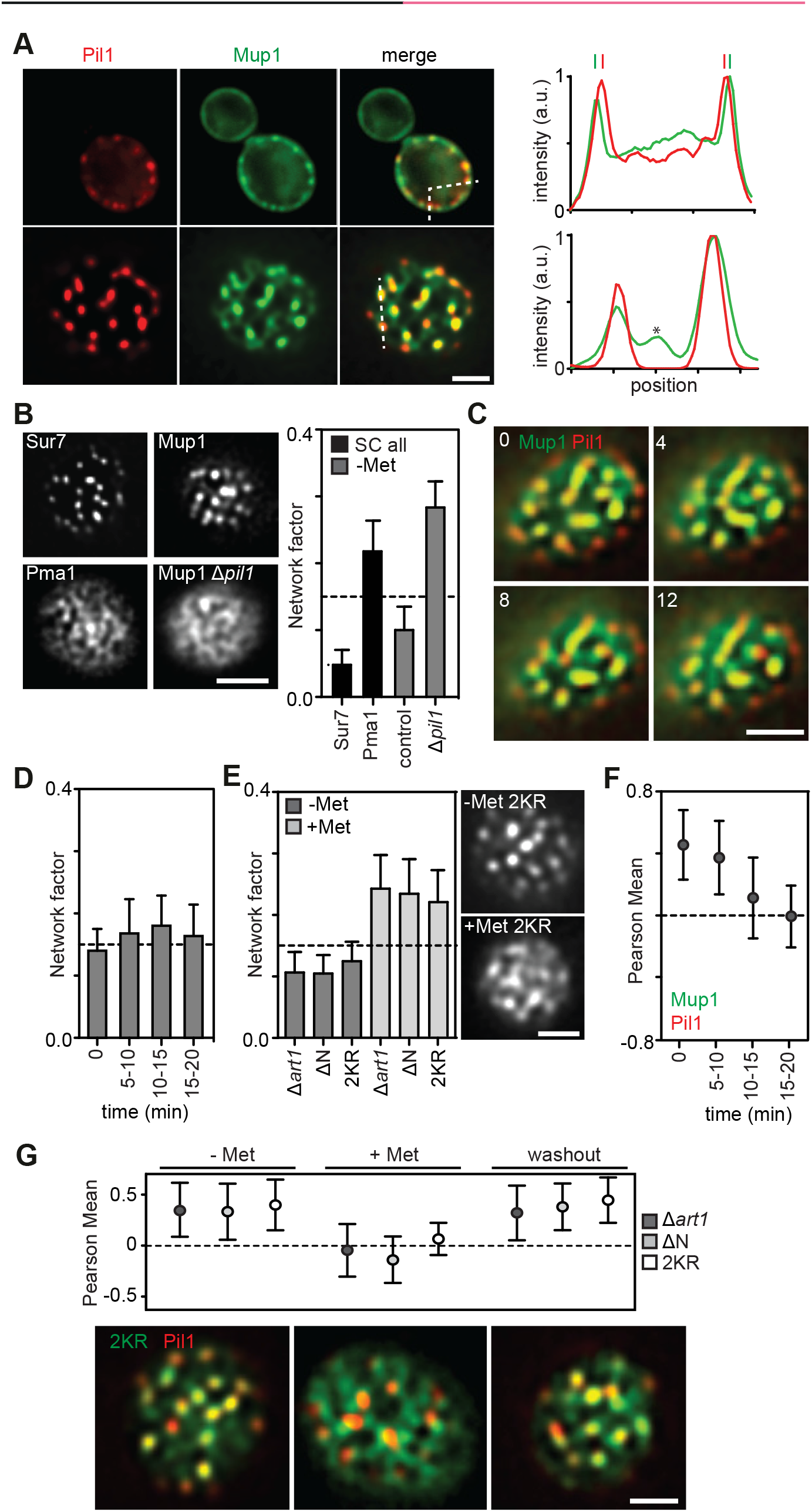
Lateral segregation of Mup1 in the yeast PM. (**A**) Equatorial and TIRFM images showing Mup1 clustering in the MCC in the absence of its substrate. Intensity profiles show Mup1 also partitioning outside the Pil1 signal (above), with additional density mapping to a network-like domain (bottom, asterisk). (**B**) Mup1-GFP intensity distribution (Network factor) indicating network (above dotted line) and patchy (below dotted line) patterns. Sur7 and Pma1 serve as references for patchy and network-like distributions, respectively. Representative TIRFM images of the different patterns are shown. (**C**) Representative images illustrate Mup1-GFP exit from the MCC labeled by Pil1-RFP upon addition of Met (timestamps in min). (**D**) Mup1-GFP distribution upon substrate addition shown by the network factor. (**E**) Mup1-GFP distribution upon substrate addition in the indicated mutants. Representative TIRFM images for the Mup1-2KR mutant are shown. (**F**) Mup1-GFP distribution upon substrate addition shown by colocalization with Pil1-RFP. (**G**) Substrate-dependent and reversible Mup1 relocation shown by quantification of its colocalization with Pil1-RFP in the indicated mutants. Representative two-color TIRFM images are shown for the Mup1-2KR mutant. Values are means ± SD, n = 50-300 cells. Scale bars: 2 μm. All values are listed in Table S1.

### Cellular regulation of Mup1 clustering

To identify cellular factors that control Mup1 clustering at the PM we investigated known regulators of MCC and eisosome integrity. Sphingolipid homeostasis has been shown to influence stability and composition of eisosomes^21–23^. In particular, Slm1 is an eisosome-resident protein that has been shown to dissociate from eisosomes and regulate TORC2 signaling upon inhibition of sphingolipid (SL) synthesis^24^. To test whether SLs also play a role in clustering of Mup1, we treated cells either with myriocin (Myr) that completely blocks SL synthesis or with aureobasidinA A (AbA), which specifically inhibits synthesis of complex SLs. As described previously, Slm1 moved out of MCC patches upon addition of either drug (Figure 3A, B). In the absence of Met, Slm1 was still able to relocate upon SL stress, albeit to a lesser degree (Figure 3A, B). Strikingly, treatment with either Myr or AbA led to pronounced relocation of Mup1 (Figure 3A, B).

**Figure 3.**
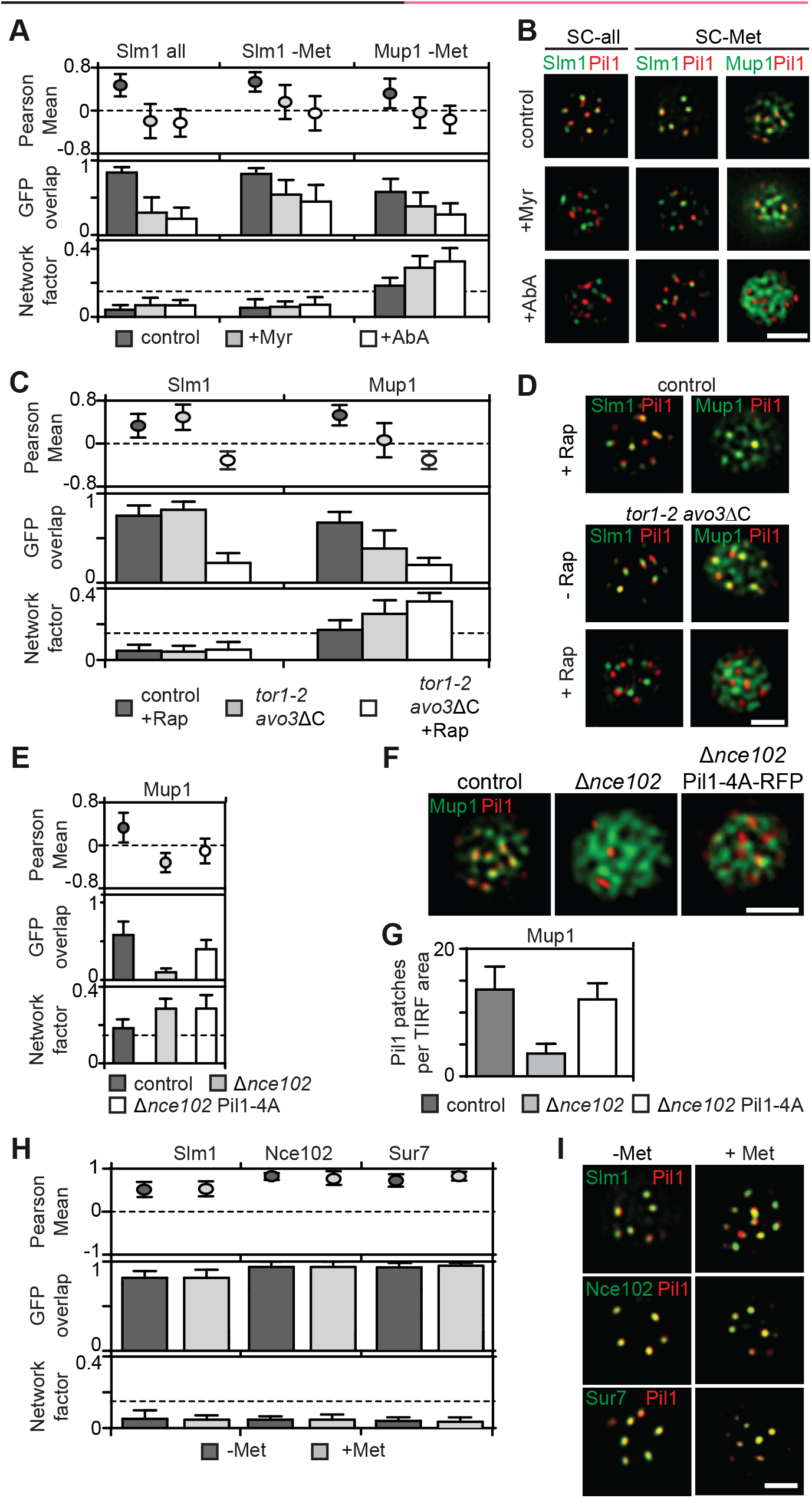
Cellular regulation of Mup1 PM clustering. (**A**) Effect of sphingolipid stress on relocation of Slm1 and on Mup1 clustering in the absence of Met. Colocalization with Pil1-RFP (Pearson Mean), GFP overlap (fraction of the GFP signal found within the structure marked by Pil1-RFP) and network factor are given. Values are means ± SD, n = 50-300 cells. Myriocin (Myr) blocks the first step in sphingolipid synthesis, while aureobasidin A (AbA) blocks the formation of complex sphingolipids. (**B**) Representative two-color TIRFM images from the experiments summarized in A. (**C**) Degrees of Mup1 clustering in the MCC in the indicated strains and upon TORC1/2 inhibition with rapamycin (Rap). The *tor1-2* mutant makes the TORC1 complex rapamycin-insensitive while the *avo3*ΔC mutant renders the TORC2 complex rapamycin-sensitive. (**D**) Representative two-color TIRFM images from the experiments summarized in C. (**E**) Nce102 is required for Mup1 clustering into the MCC. (**F**) Representative two-color TIRFM images of the Nce102-dependent clustering of Mup1-GFP in the PM from the experiments summarized in E. (**G**) Nce102-dependent eisosome stability shown as number of Pil1-RFP or Pil1-4A-RFP patches per area. (**H**) Lateral segregation of Slm1, Nce102 and Sur7 before and after Met addition. (**I**) Representative two-color TIRFM images from the experiments summarized in H. All values are means ± SD, n = 50-300 cells. Scale bar: 2 μm. All values are listed in Table S1.

As Slm1 has been implicated in the regulation of the TORC2, we next asked whether TORC signaling is involved in Mup1 clustering. We generated strains in which either TORC1, TORC2, or both could be inhibited by the addition of the drug rapamycin^25^ (Rap). As expected from its role in protein synthesis and sorting^26^, TORC1 but not TORC2, was found to be required for delivery of Mup1 to the PM (Figure S2A). Conversely, and in agreement with the reported function of TORC2 in actin organization and endocytosis^27^, inhibition of TORC2, but not TORC1, blocked internalization of Mup1 (Figure S2B). We found that inhibition of TORC2 led to the release of Slm1 from the MCC within 30 min (Figure 3C, D). Importantly, Mup1 also dissociated from the MCC upon inhibition of TORC2, but not of TORC1 (Figure 3C, D). Reduced Mup1 clustering already became apparent when simply removing the C-terminus of the TORC2 subunit Avo3 (to render the complex sensitive to Rap, Figure 3C), indicating that this mutation partly affects TORC2 function and MCC integrity. In summary, our results show that both SLs and TORC2 signaling regulate clustering of Mup1 in the MCC.

While TORC2 inhibition induced full Mup1 relocation within 15-30 minutes, Myr and AbA treatment had to be maintained for more than 1 h to achieve comparable results. We therefore wanted to test whether TORC2 could also regulate Mup1 segregation through additional factors and tested the localization of other MCC components in our conditions. The tetraspanner protein Nce102 has been shown to support the segregation of several transporters into the MCC^9^ and to exit MCC patches upon SL perturbation^28^. In agreement with this report, we found that Nce102 partly moved from MCC patches into a network-like compartment upon treatment with Myr and AbA in rich medium (SC all, Figure S2C, D). Importantly, neither Nce102 nor another MCC-resident tetraspanner, Sur7, showed significant relocation when SL synthesis was blocked in the absence of Met (Figure S2C, D). When we inhibited TORC2 signaling in cells grown in the absence of Met, we found that Sur7 localization was again unaffected. However, Nce102 was largely lost from the MCC and was dispersed all over the PM (Figure S2E, F). To test whether this effect contributes to the reduced Mup1 clustering upon TORC2 inhibition, we deleted *nce102*. Strikingly, Mup1 was almost completely lost from the MCC in Δ*nce102* cells (Figure 3E, F). A known additional phenotype of *nce102* deletion is the disassembly of eisosomes through Pkh-dependent phosphorylation of Pil1^28^. To prevent this effect we expressed a non-phosphorylatable mutant of Pil1 (Pil1-4A). Even though the number of eisosomes was restored in this mutant (Figure 3G), Mup1 remained fully dispersed in the PM (Figure 3E, F). In summary, our results indicate that the tetraspanner Nce102 is essential but not sufficient to ensure clustering of Mup1 into the MCC.

Importantly, the effects of SL and TORC2 inhibition on Mup1 clustering occurred within 30-60 min, while relocation of Mup1 upon Met addition must occur within a few minutes (Figure 1, 2). In addition, all potential regulators of Mup1 clustering, Sur7, Nce102 and Slm1, did not become relocated out of MCC clusters by addition of Met (Figure 3H, I). We therefore investigated whether intrinsic features of Mup1 directly controlled its rapid lateral relocation within the PM.

### PM relocation of Mup1 is controlled by a conformational switch

Neither clustering to the MCC nor relocation of Mup1 required ubiquitination or the cytosolic N-terminal segment of Mup1. We therefore focused our analysis on intrinsic features of Mup1 that might contribute to its localization and dynamics on the C-terminal tail and the transmembrane segments of the permease. In transporters that belong to the Amino acid-Polyamine-organoCation (APC) superfamily, including Can1^19^ and Fur4^29^, substrate uptake has been proposed to induce a structural transition from an open to a closed conformation. Based on crystal structures of several bacterial APC transporters, Mup1 is expected to adopt a 5+5 fold structure with two inverted repeats of 5 transmembrane domains (TMDs)^30^. Substrate binding in this fold is stabilized by glycine-rich regions within TMDs 1 and 6, and a stack of aromatic side-chains within TMDs 3, 6 and 10^19^. We mutated glycine 78 (G78N) in TMD1 and tryptophan 155 (W155A) in TMD3 to test the role of these structural features in Met transport by Mup1. Both residues are conserved among fungal transporters and the glycine also occurs in bacterial homologues (Figure S3A). Based on a high-confidence structural homology model (Figure S3B), they are expected to extend into the substrate-binding pocket. In accordance with that prediction, mutation of the conserved glycine completely inhibited Met uptake, and replacement of the aromatic tryptophan by an alanine residue led to a marked reduction in Met transport (Figure 4A). However, while the G78N mutant is resistant to the toxic Met analog selenomethionine (SeMet, Figure 4B), the W155A substitution actually enhances sensitivity to SeMet. These observations can be explained by the reductions in Mup1 ubiquitination and endocytosis seen in both mutants (Figure 4C-E). The expression of PM-stabilized (due to impaired ubiquitination) Mup1 variants with full or reduced function (W155A, ΔN) should lead to higher Met (or SeMet) accumulation, whereas lack of ubiquitination of the completely inactive G78N variant in the PM would have no effect on sensitivity to SeMet. Note that the reduced Met uptake by Mup1-ΔN (Figure 4A) is most probably attributable to its partial retention at the ER, and its correspondingly lower expression at the PM (Figure 4D, Figure S3C). In contrast to the endocytosis-deficient mutants introduced above (ΔN, 2KR; see Figure 1F), G78N – and to a lesser extent W155A – exhibited reduced exit from the MCC region upon Met addition, as measured by an increase in the network factor (Figure 4F) and an accompanying reduction of colocalization with Pil1-RFP (Figure 4G). These results indicate that the conformational switch associated with substrate transport is required for both lateral relocation and endocytic uptake of Mup1.

**Figure 4.**
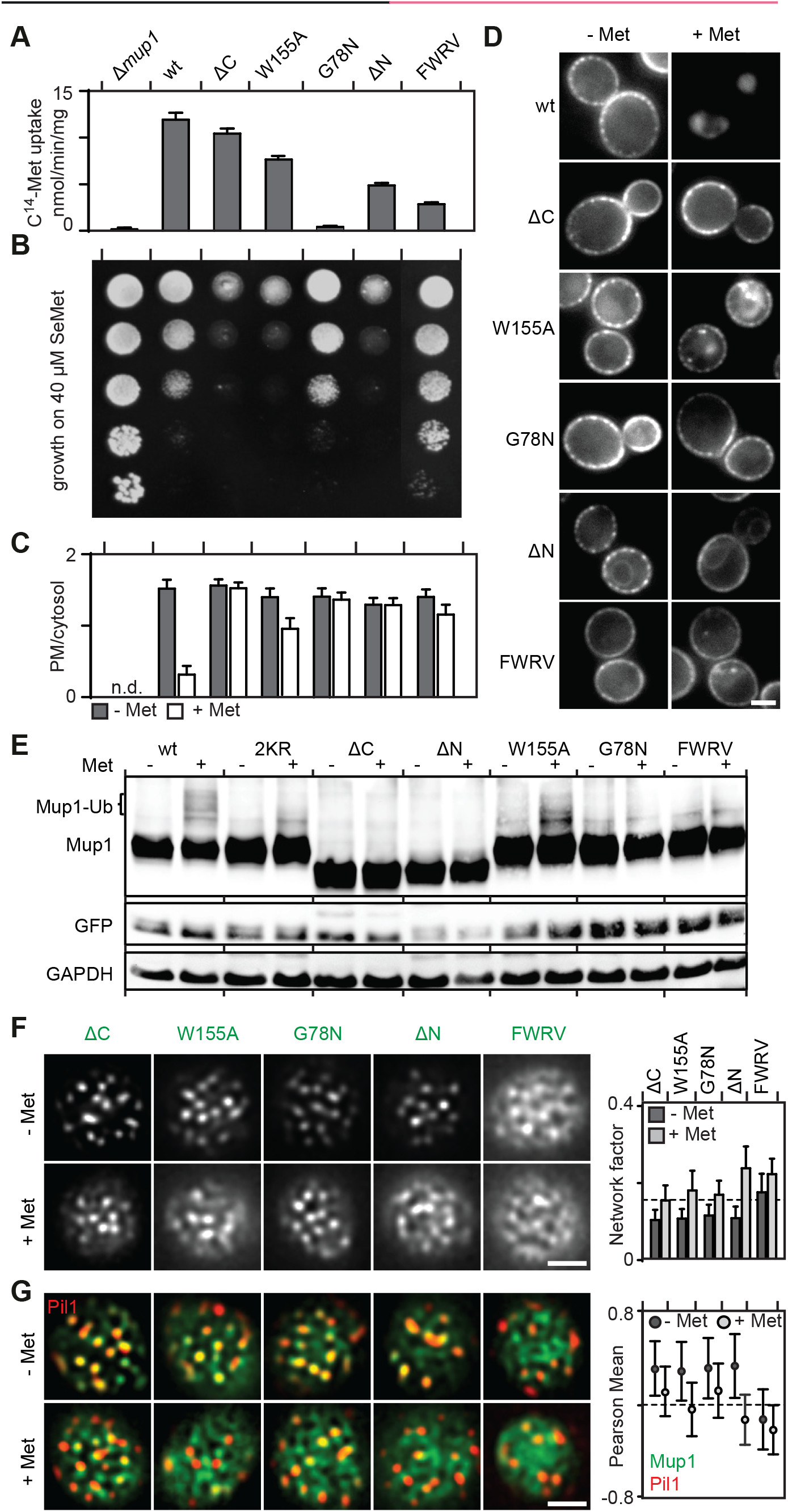
A conformational switch drives Mup1 domain relocation at the PM. (**A, B**) Comparisons of the functional activities of different Mup1 mutants based on direct quantification of ^14^C-Met uptake (**A**) and growth sensitivity to the toxic Met analog selenomethionine (SeMet, **B**). (**C, D**) Endocytic internalization of the different Mup1 mutants fused to GFP. Ratios of PM to cytosolic fluorescence intensities (**C**) and representative images at equatorial planes (**D**) are shown. (**E**) Western blot showing ubiquitination (Ub) of the indicated Mup1 mutants grown in the absence (-) and presence of Met (+, 5 min after addition). Loading control as in Figure 1C. (**F, G**) Lateral PM segregation is shown in representative TIRFM images and quantified in terms of the network factor (**F**) or the degree of colocalization with Pil1-RFP (**G**). Mutants utilized: wt, wild type Mup1; ΔC, deletion of the segment C-terminal to aa519; W155A and G78N, point mutants; ΔN, deletion of the region N-terminal to aa52; FWRV, mutation of aa534-537 to alanines; 2KR, Mup1 with N-terminal mutations K27R and K28R. All strains refer to Mup1 variants fused to C-terminal GFP. Values are means ± SD, n = 2-4 experiments (A) and n = 50-250 cells (C, F, G). n.d.: not determined. Scale bars: 2 μm. All measured values are listed in Table S1.

As substrate binding was not required for Mup1’s ability to cluster into the MCC (Figure 4F), we turned to an analysis of the cytosolic C-terminus of Mup1. When we removed the C-terminus (ΔC = aa520-end), Mup1’s transport activity was largely unchanged (Figure 4A), but the protein was no longer ubiquitinated and remained at the PM upon addition of Met (Figure 4C-E). As expected, this led to hypersensitivity to SeMet (Figure 4B, Figure S4). In addition, while clustering into MCC patches was unaffected (as in the case of the N-terminal truncation mutant), exit from the MCC domain was compromised in cells expressing ΔC (Figure 4F, G). Complete deletion of both N-and C-terminus did not further exacerbate the phenotype (Figure S3C, D). The C-terminal segment of Mup1 has multiple potential phosphorylation and ubiquitination sites (Figure S3E). However, removal of these sites (in mutants Δ543 and Δ565) did not have any noticeable effects on Mup1 function or internalization (Figure S4), suggesting that C-terminal post-translational modifications do not contribute to the lateral segregation and the endocytic turnover of Mup1. When comparing Mup1 with known crystal structures of APC transporters, we noticed a structural domain near the beginning of the exposed C-terminal segment (positions 512-543) that mimics a predicted loop in the bacterial GadC transporter with high confidence (Figure S3B, E). Interestingly, this loop was identified as a C-terminal “plug” that reaches into the core of the TMD segments in GadC^31^. In Mup1 the tip of this plug should be positioned close to the substrate-binding pocket and channel (Figure S3B). Removal of the C-terminal sequence of the plug up to position 543 did not affect either the function or internalization of Mup1 (Figure S4A-D). However, further truncation by an additional 6 or 10 amino acids (Δ537 and Δ533) led to phenotypes similar to those seen upon complete deletion (Δ520) of the C-terminal segment in the mutant ΔC (Figure S4). The segment most relevant for Mup1 localization and function (FWRV, positions 534-537) corresponds to a conserved motif found within the C-terminal regions of fungal Mup1 homologues (Figure S3E, F). To test for a possible role of this signature sequence in Mup1 domain localization, we mutated all four amino acids to alanines (to yield the ‘FWRV’ mutant). In contrast to the full C-terminal truncation, the FWRV Mup1 mutant exhibited strongly reduced function (Figure 4A, B) but was similarly defective in ubiquitination and internalization (Figure 4C-E). This is consistent with the previously proposed role of the C-plug in facilitating the conformational transition during substrate transport in GadC^31^. Strikingly, we found that the FWRV mutant was no longer able to cluster into the MCC domain in the absence of Met, and was instead distributed in a network-like fashion (Figure 4F, G). Moreover, addition of Met had only a mild effect on this pattern. This is consistent with the FWRV mutant adopting a closed conformation that is usually occupied only transiently during the substrate transport cycle. In summary, our data support a role for the conformational state of Mup1 in mediating its clustering in the MCC and its rapid lateral relocation upon substrate transport. Our results also suggest a critical role of the C-terminal plug in the conformational switch.

### Mup1 relocates into a unique dispersed network

We have previously shown that the yeast PM consists of a multitude of distinct domains^4^. So far we have discussed how Mup1 clusters in MCC patches, but we have not defined the region of the PM into which it relocates upon its release from the MCC. To better define the network-like domain where Mup1 resides after substrate addition (Figure 2), we tested its colocalization with the MCP marker Pma1 (Figure 5A), which is often used as a general marker for all PM areas outside of the MCC^20^. Interestingly, while we confirmed the perfect separation of the MCC and the MCP (Figure 5A), we found that even after MCC exit Mup1 clearly segregated from Pma1 (Figure 5A). This segregation was as strong as that between the MCC and the MCP (Figure 5A, right), indicating that Mup1 localizes to a distinct network domain upon MCC exit. As the limited resolution of our TIRFM images does not allow precise definition of the areas covered by particular proteins in the PM, we further characterized the different network domains via 3D structured illumination microscopy (3D-SIM). This approach confirmed mutual segregation of the three domains (Figure 5B).

**Figure 5.**
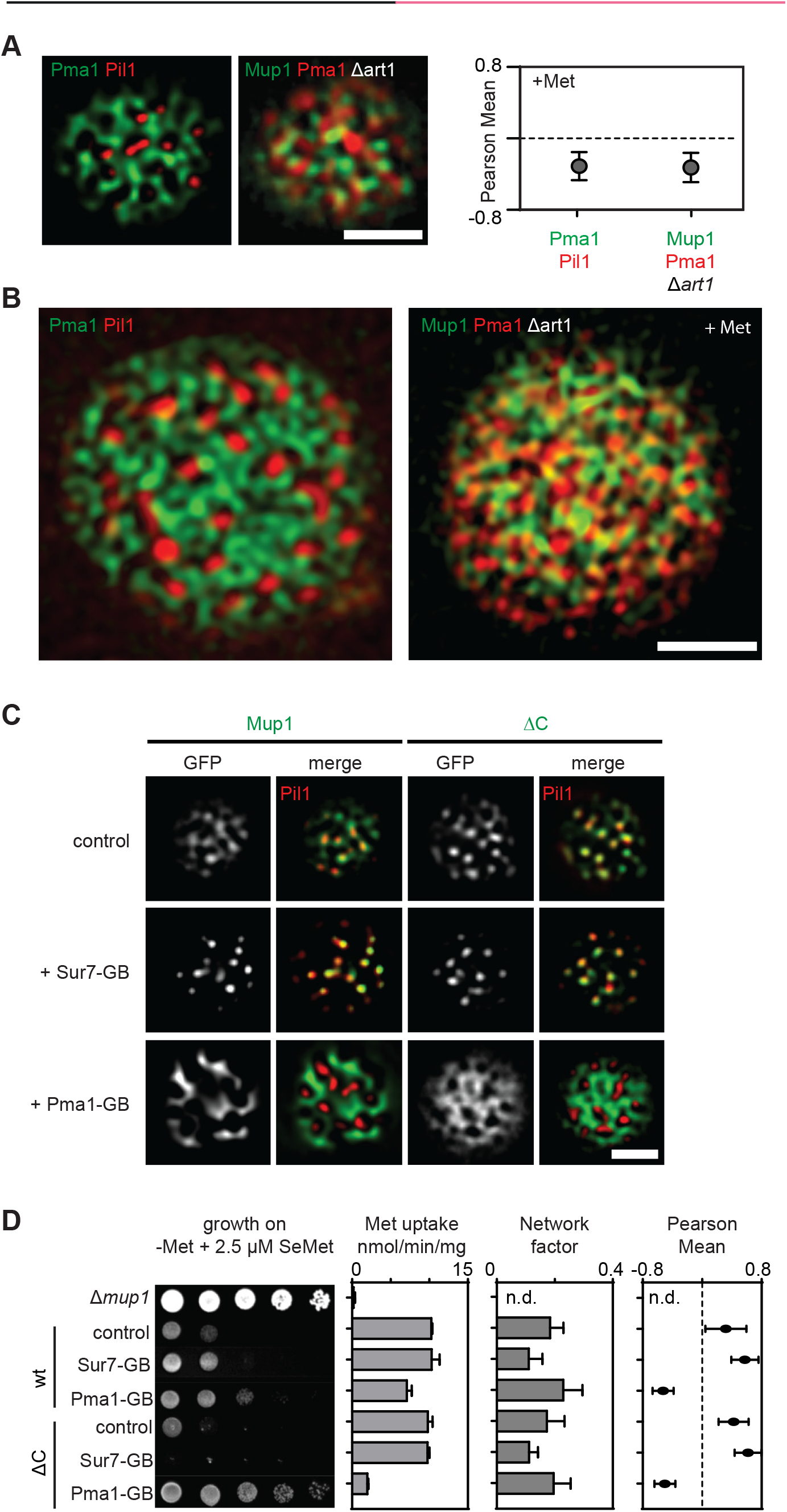
Spatial segregation of Mup1 into a unique dispersed network upon MCC exit. (**A**) Representative two-color TIRFM images showing the spatial separation between the MCC (shown as Pil1-RFP) and the MCP (shown as Pma1-GFP), and the distribution of Mup1 outside the MCC together with the MCP marker Pma1-RFP in the absence of ubiquitination and upon Met addition. The graph shows the quantification of colocalization (Person Mean) for the indicated protein combinations. (**B**) Representative 3D-SIM images showing the distribution of the indicated protein pairs in A. (**C**) TIRFM images showing Mup1 and ΔC artificially recruited to different PM domains using the GFP trap system. GB, GFP-binding nanobody. Pil1-RFP was used to illustrate the degree of relocation out of the MCC. (**D**) Effects of forced relocation on SeMet sensitivity, rate of Met uptake and the distribution of the respective Mup1 variant (Mup1/ΔC) in the PM (n.d., not determined). Note that the used concentration of SeMet (no added Met) was not sufficient to induce endocytosis of Mup1 (not shown). Values are means ± SD, n = 2-4 experiments (uptake), n = 50-200 cells (Network factor and Pearson Mean). Scale bars: 2 μm. All measured values are listed in Table S1.

To determine whether the spatial segregation into the Mup1 network domain is relevant for its activity, we artificially relocated Mup1 to various domains using the GFP trap system (Figure 5C). Strikingly, we found that relocation of Mup1 to the Pma1 domain reduced its biochemical activity (Figure 5D). This effect was seen with the WT variant, but was much more striking for the ΔC mutant (Figure 5D). Pma1 function was not affected by fusion to the GFP trap (Figure S5A). The reduction in Mup1 activity was also not due to a general requirement of the MCC environment, as deletion of *pil1* (with concomitant loss of eisosomes) did not reduce Met uptake activity by Mup1 (Figure S5B). Moreover, forced attachment of Mup1 to Sur7 did not noticeably increase Met uptake (Figure 5D). In contrast, the arginine transporter Can1 exhibited increased activity and sensitivity to canavanine (a toxic analog of arginine) when anchored to the MCC (Figure S5C).

In summary, we found that upon MCC exit Mup1 segregates into a unique network-like domain. Of particular interest is the observation that the proton pump Pma1, or the PM environment of the MCP inhibit Mup1 activity. This might be a general effect on proton symporters, as we observed a similar, but even more pronounced effect on canavanine uptake via Can1 (Figure S5C).

### Mup1 clusters are protected from endocytosis

The MCC and eisosomal PM region has been proposed to be refractory to endocytosis^9^, although this hypothesis has since been challenged^32^. To test whether accumulation of Mup1 in the MCC hinders its endocytic re-uptake – or simply precludes the prerequisite ubiquitination step – we artificially fused Mup1 to a single ubiquitin (Ub) moiety^33^. As expected, Mup1-Ub was constitutively localized to the vacuole, even in the absence of its substrate (Figure 6A). In contrast, the ΔC mutant, which is defective in MCC exit, was partially retained in clusters at the PM (Figure 6A). Importantly, disruption of MCC domains by deletion of *pil1* completely abrogated ΔC-Ub retention at the PM (Figure 6A). This is consistent with the idea that occupants of MCC domains are protected from endocytosis. As ΔC without fused Ub was retained at the PM in a Δ*pil1* strain grown with Met, our data also suggest that the C-terminal plug is directly involved in Mup1 ubiquitination (Figure 6B). As a further test of the effect of the MCC on endocytosis, we attached Mup1 directly to the abundant MCC marker Nce102 (Figure 1B) using the GFP trap system. As expected, anchoring of Mup1 to the MCC protected the permease from Met-dependent degradation, and this effect was abolished when *pil1* was deleted (Figure 6C). Finally, when we followed the effects of endocytic protection in yeast cultures grown for longer periods in the absence of Met, we observed slow internalization of Mup1 at a rate that was significantly enhanced in the absence of *pil1* (Figure 6D). In summary, our data are consistent with a role of the MCC domain in protecting resident proteins from endocytic uptake and degradation.

**Figure 6.**
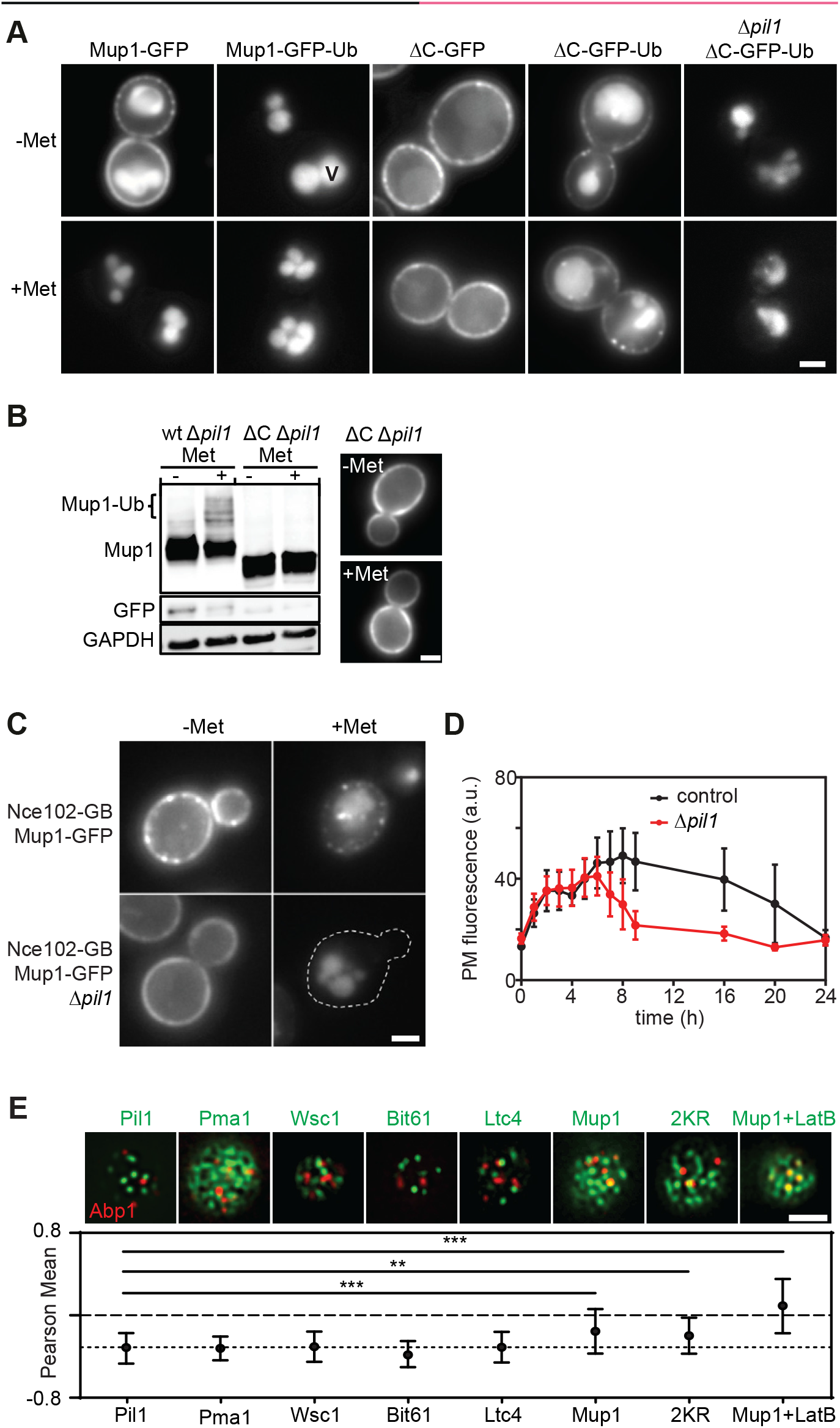
Clustering of Mup1 protects from endocytosis. (**A**) Influence of the in-frame fusion of a ubiquitin (Ub) moiety to Mup1 and ΔC on PM localization and turnover. V = vacuole. (**B**) Western blot showing Mup1 and ΔC ubiquitination in Δ*pil1*. Samples were prepared 5 min after addition of Met. Loading control as in Figure 1C. Representative images of indicated conditions are shown to illustrate defect in endocytosis of ΔC. (**C**) Artificial anchoring of Mup1 to Nce102 in the presence or absence (Δ*pil1*) of MCC and Met, respectively. (**D**) Mup1 internalization during long-term growth in the absence of Met. (**E**) Degree of colocalization of endocytic events (shown by the late endocytic marker Abp1-RFP) with various PM domains (GFP fusions). Representative two-color TIRFM images of the indicated protein pairs are shown. Pil1 represents eisosomes, Pma1 the MCP, Wsc1 the MCW (membrane compartment occupied by Wsc1), Bit61 the MCT and Ltc4 the MCL (membrane compartment of Ltc3/4). Cells were grown in Sc-Met and imaging performed after addition of 100μM Met. LatB was used at a concentration of 100 μM to stabilize endocytic sites. The dotted line is used as a guide to the eye. Asterisks indicate significantly different data sets: ** p<0.005, *** p<0.001. Values represent means ± SD, n = 50-150 (D) n = 25-80 (E). Scale bars: 2 μm. All measured values are listed in Table S1.

In light of our finding that Mup1 occupies a distinct domain in the PM even after exit from the MCC, we wondered whether endocytosis might be specifically facilitated by this network domain. To test for a preferred association of Mup1 and endocytic sites we colocalized various integral PM markers with the late endocytic marker Abp1-RFP (Figure 6E). As expected from the observed protection from turnover in MCC patches, endocytic patches were completely segregated from Pil1 (Figure 6E). Indeed, we found that all of the previously described domains clearly segregated from endocytic regions (Figure 6E). Interestingly, 15 minutes after Met addition Mup1 or its 2KR variant were significantly less segregated from endocytic regions than all other tested markers (Figure 6E). This association was strongly enhanced when endocytic sites were stabilized by addition of low doses of Latrunculin B (LatB, Figure 6E). Our results thus show that the Mup1 molecules in a specific PM network outside of MCC clusters partially overlap with endocytic sites. Given that Mup1 becomes internalized within 10 minutes of relocating from the MCC (Figure 1D), this raises the question, whether Mup1 can directly influence distribution of endocytic sites.

### Mup1 actively redirects endocytic events

To test this hypothesis we tracked the late endocytic marker Abp1, together with Mup1, at different time points after Met addition to monitor Mup1 endocytosis. We found no significant colocalization of actin patches with Mup1 in the initial phase of Mup1 relocation (not shown). However, within 10 min of Met addition, multiple Abp1 patches assembled on or close to Mup1 clusters (Figure 7A, Movie S1, asterisks). Importantly, in most of these cases, both markers disappeared or were depleted together (Figure 7A, line scans shown for two examples). When we blocked internalization of actin patches by treatment with 100 μM LatB, we observed extended colocalization of Abp1 with Mup1 clusters (Figure 7B). In addition, upon disassembly of Abp1 patches, the Mup1 signal remained unchanged, which is consistent with the observed lack of internalization within 1 h of Met addition (not shown). These results indicate that clusters of ubiquitinated Mup1 can act as instructive signals for the recruitment of endocytic sites. To verify this we quantified the distribution of endocytic patches in polarized yeast cells with small-to medium-sized buds. In these cells, most endocytic events are limited to the bud-region, where all polarized growth occurs. Remarkably, within 2-3 min of Met addition, we observed a large increase in endocytic activity within mother cells, with a coincident decline in buds (Figure 7C). This ‘depolarization’ of endocytosis preceded the observed uptake of Mup1 into cells (Figure 7C, right panel). When we followed this response over an extended period we found that the depolarization reversed within 10 minutes and all endocytic patches returned to the buds within 20-30 min (Figure 7D). Importantly, the response was not observed for cells that either did not express any Mup1 or that only expressed the ubiquitination-deficient 2KR mutant. Together, our results are consistent with an active role for ubiquitinated Mup1 in driving its own internalization and regulating the subcellular distribution of endocytosis.

**Figure 7.**
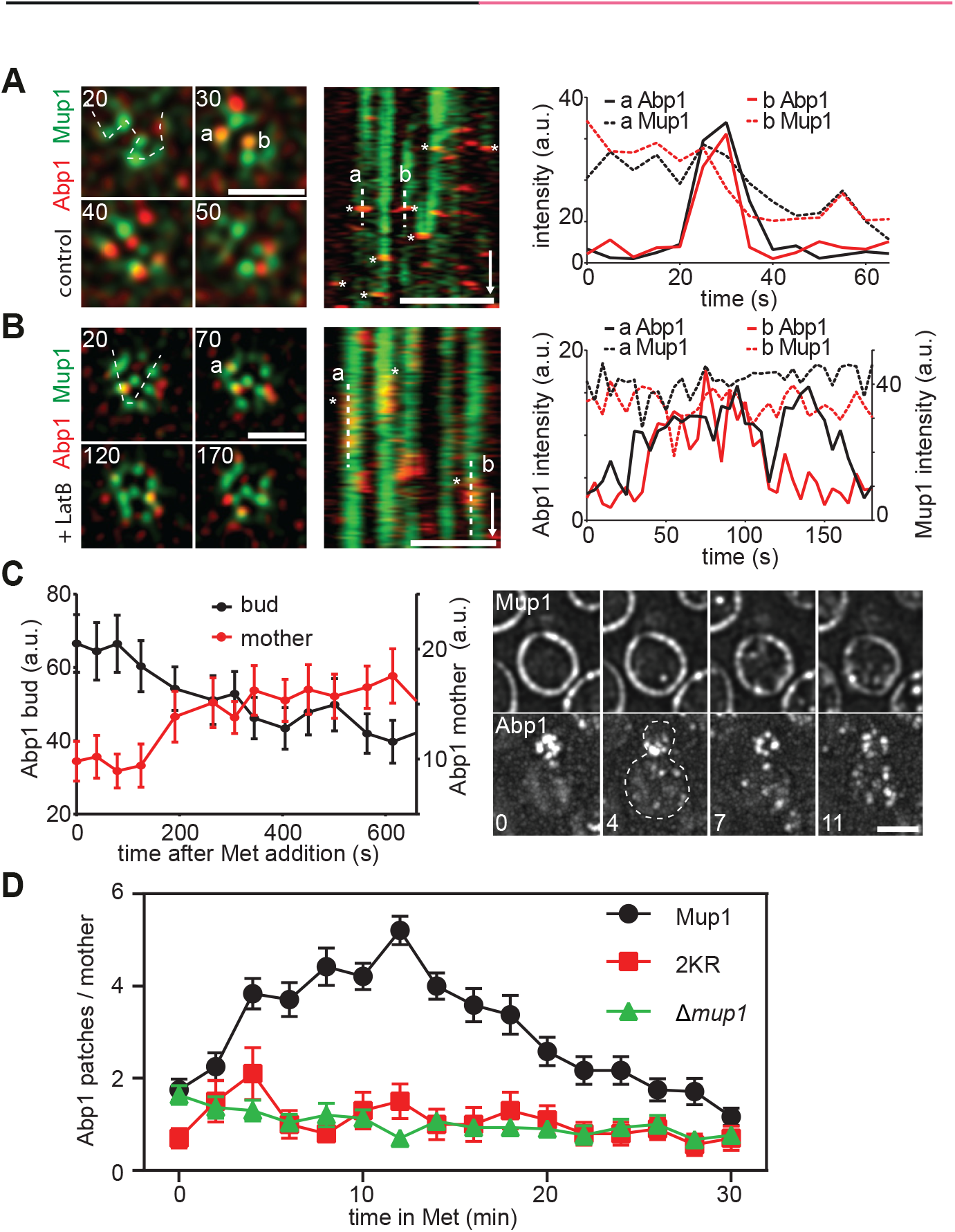
Mup1 actively redirects endocytic events. (**A**) Correlation of endocytic patch localization (marked by Abp1-RFP) with Mup1 distribution. Kymographs (along the dotted line in the indicated cells) show localization patterns starting 10-15 min after Met addition. Representative line scans illustrate coordinated assembly and removal of endocytic patches from the indicated traces in the kymographs (marked “a” and “b”). Timestamps are given in s, time arrow: 100 s. (**B**) Similar to the experiment in A, except for the presence of 100 μM LatB. Note the longer lifetime of endocytic patches and the fact that there is no change in the fluorescence of Mup1-GFP upon disassembly of Abp1-RFP. (**C**) Changes in the relative intensity of the Abp1-RFP signals in the bud (black) and the mother cell (red) following the addition of Met. Representative images of Abp1-RFP and Mup1-GFP of a polarized cell are shown. Timestamps are given in min. (**D**) Number of Abp1-RFP patches in the mother cell during Met addition in the indicated strains. Growing cells with small buds and highly polarized Abp1 were individually selected, and the amount of Abp1 patches in the mother cell manually quantified. Values represent means ± SEM, n = 18-24 (C), n = 10-30 (D) cells. Scale bars: 2 μm. All measured values are listed in Table S1.

## Discussion

Cells must adapt rapidly to a wide variety of environmental changes and stresses. One universal mechanism that facilitates such adaptation is the selective alteration of the array of nutrient transporters present at the PM^34^. In budding yeast, 20-60% of energy consumption in exponentially growing cells is devoted to the maintenance of a proton gradient across the PM^35^, which serves as the primary driving force for the activity of a wide range of nutrient transporters. Yeast cells therefore face the critical challenge of providing optimal PM levels of nutrient transporters that ensure adequate nutrient uptake, while avoiding unnecessary energy consumption. Here we provide evidence that controlled lateral segregation of transporters within specific PM domains provides a novel and important level of regulation in the response of cells to nutrient availability. We suggest that lateral segregation of the PM contributes to an optimal utilization of energy and resources.

The high-affinity methionine permease Mup1 constitutes an ideal model with which to study the mechanisms of nutrient adaptation. Induction of Mup1 expression occurs promptly upon removal of its substrate, and Mup1 levels in the PM quickly surpass those of every other integral PM protein. Conversely, addition of even micromolar levels of substrate leads within minutes to endocytic uptake and degradation of Mup1. Importantly, we found that, as a prerequisite to its rapid turnover, Mup1 is also regulated via lateral segregation within the PM. In the absence of its substrate, Mup1 is clustered in the patchy MCC domain, where it is protected from internalization. Upon substrate addition, however, the transporter quickly leaves the MCC and relocates to a defined network-like domain (Figure 8A). This relocation does not depend on ubiquitination but is inhibited by mutations that block substrate binding. We also uncovered a critical role for a C-terminal domain of Mup1 in mediating lateral relocation. This domain is similar to one which is predicted to insert into the core of the structurally homologous bacterial transporter GadC^31^. Deletion of the “C-plug” (ΔC) in Mup1 did not interfere with substrate transport, but completely inhibited its lateral relocation, ubiquitination and subsequent internalization. We found that ubiquitination of ΔC was blocked in cells, even when the MCC had been disrupted by depletion of *pil1*. In addition, a mutant with alterations in the tip of the C-plug (FWRV→AAAA) exhibited strongly reduced function and was no longer able to cluster into the MCC. This mutant was also not susceptible to ubiquitination and exhibited very little substrate-induced internalization. Our results therefore indicate that the C-plug plays a structural role in Art1-mediated ubiquitination of the Mup1 N-terminus and is critical for efficient lateral relocation. Interestingly, a recent study reported a negatively charged region in the membrane-proximal part of the N-terminal tail (residues 40-55) of Mup1 as interaction site for Art1^18^. However, this binding site was not sufficient for Mup1 ubiquitination and a second interaction site was postulated. We propose that the C-plug serves as a linker domain that transmits the conformational change during substrate transport to the cytosolic N-terminal tail, thereby exposing the binding site for Art1.

**Figure 8.**
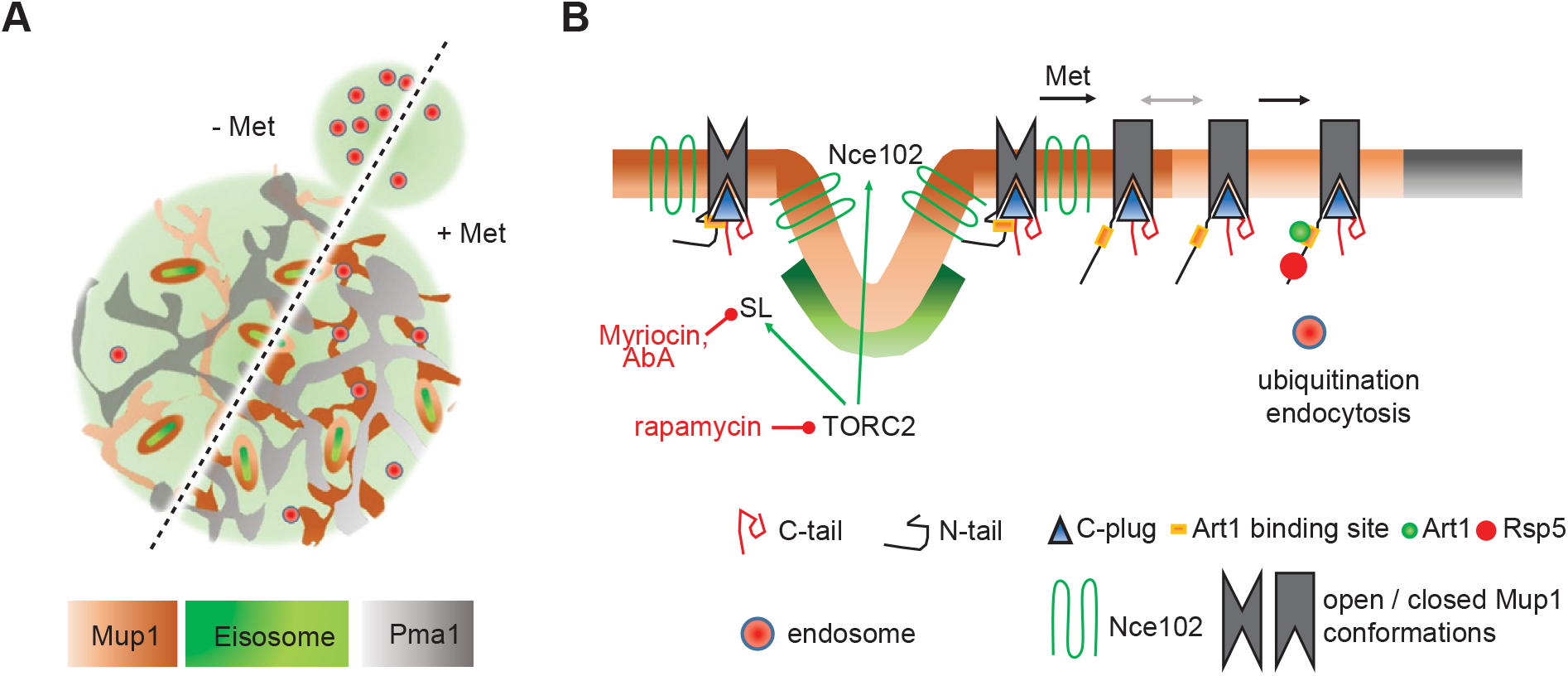
Model for the regulation of Mup1 segregation and turnover. (**A**) Lateral organization of the yeast PM indicating relevant domains occupied by the indicated proteins. Upon addition of its substrate (+ Met), Mup1 relocates from the MCC (brown area around eisosomes) into a network-like domain (light or dark brown), which is distinct from that marked by Pma1 (grey). Endocytosis in polarized cells is usually focused in the bud region but becomes depolarized upon addition of methionine. (**B**) During substrate transport Mup1 undergoes a conformational change (indicated by the different grey shapes) that facilitates exposure of the N-terminal ubiquitination sites and exit from the MCC (grey arrow). Ubiquitination and domain relocation both require an intact C-plug (blue triangle). Exposure of the N-terminus permits binding of Art1 (green circle) and subsequent recruitment of the ubiquitin ligase Rsp5 (red circle). Once modified, Ub-Mup1 recruits the endocytic machinery (red circle with blue rim) and becomes internalized. The legend describes all indicated domains and proteins.

We show here that clustering of Mup1 into MCC patches is regulated by SLs, by the MCC-localized tetraspanner Nce102 and by TORC2 signaling (Figure 8B). Our data indicates that a complete blockade of SL synthesis (addition of Myr) or specific inhibition of the production of complex SLs (treatment with AbA) interferes with Mup1 clustering. SL synthesis is under control of the central metabolic regulatory kinases TORC1 and TORC2^36^. Strikingly, we found that selective inhibition of TORC2 but not TORC1 abolished clustering of Mup1 in the MCC. In addition to regulating SL levels, we found that TORC2 activity was also required for MCC localization of the tetraspanner protein Nce102, which is even more abundant in MCC patches than the core protein Pil1. Interestingly, Nce102 was not lost from the MCC following treatment with Myr or AbA. This contradicts previous findings^28^ and might be attributed to a distinctive mode of regulation in the absence of Met. Importantly, deletion of *nce102* led to a complete loss of Mup1 from the MCC, and this effect could not be reversed by stabilization of eisosomes via expression of non-phosphorylatable Pil1. Our results thus indicate that TORC2 regulates Mup1 clustering via two distinct mechanisms – SL biosynthesis and Nce102. SL stress could alter the global biophysical properties of the PM, thereby affecting composition and organization of individual domains. Considering the patchwork nature of the yeast PM^4^, we favor a model where cooperative weak interactions between proteins (Nce102, Mup1) and lipids (SL) can drive stable lateral segregation within the plane of the bilayer. Irrespective of the actual regulators, we show that conformational changes within Mup1 are sufficient to rapidly override the existing lateral interactions and thereby mediate rapid lateral relocation of the permease. Our findings are consistent with a dynamic regulation of lateral segregation that is highly adaptable to the environment and metabolic requirements of the cell.

In keeping with our previous report on the existence of a multitude of PM domains in yeast ^4^, the network-like domain defined by Mup1 after its exit from MCC patches is clearly distinct from the previously defined MCC and MCP domains. We found that forced anchoring of Mup1 to the proton pump Pma1 reduced its catalytic activity. This effect was even more apparent for the ΔC mutant, which lost nearly all its activity upon relocation. For GadC it has been shown that the compact C-plug covers a cleft in the protein core that has many exposed charges^31^. In addition, removal of the C-plug altered the pH sensitivity of the transporter^31^. It is therefore tempting to speculate that the Pma1 domain constitutes a physicochemical environment that is harmful for Mup1. Since anchoring of the arginine transporter Can1 to Pma1 also abolishes its transport activity^4^, this effect of the Pma1 compartment appears to be equally valid for other proton symporters.

Finally, we found that lateral segregation of Mup1 directly regulates its endocytic turnover. It has previously been shown that endocytosis is excluded from MCC patches^9, 32^, and we confirmed that Mup1 is not internalized while attached to the MCC. Importantly, we now show that endocytic internalization preferentially occurs at sites of Mup1 accumulation after substrate-induced relocation from the MCC and ubiquitination. We specifically demonstrate for polarized cells that, within minutes of Met addition, endocytic events are redirected from the bud to the network of Mup1 in mother cells. Our results therefore indicate a dual role for Mup1 segregation: On the one hand the MCC acts as protective site that ensures retention of functional Mup1 at the PM. On the other hand, ubiquitinated Mup1 that exits from the MCC acts as a potent beacon for endocytic adaptor proteins and is therefore able to efficiently compete with the bud-localized signals that drive endocytosis to polarized growth regions.

Our study demonstrates for the first time that a transition between different types of lateral domains can regulate protein function and turnover at the plasma membrane. We not only elucidate the mechanism for lateral segregation of the Mup1 permease but also suggest a specific biological function for this process. Lateral segregation is a fundamental property of all biological membranes. It is therefore tempting to speculate that the uncovered interplay between protein segregation and turnover will apply to many other membrane components and cell type.

## Methods

### Yeast strains and plasmids

All strains created in this study were derived from the *Saccharomyces cerevisiae* BY4741 Mat**a** strain (Euroscarf). In addition, strains from the UCSF C-terminal GFP-fusion collection^37^, and EUROFAN II KO collection ^38^ were used. Genomic carboxy-terminal tagging and gene deletions were performed by direct integration of PCR products as described previously^39^. All mutants based on the Δ*mup1* strain were generated by replacing the kanMX KO cassette with the required PCR product (Mup1mutant-tag-hphNT1) through *in vivo* homologous recombination. The PCR product was generated through overlapping PCR using standard methods. The *tor1-2* mutant was generated by homologous recombination with a 100 bp long oligonucleotide containing the required mutation^40^ and selection on plates containing 1 μg/ml rapamycin. Plasmids were constructed using standard molecular biology techniques. Transformation into yeast cells was performed using the LiOAc method. All plasmids were sequence-verified. All PCR-derived endogenous integrations were verified via colony-PCR and sequencing. All strains and plasmid used in this study are listed in Table S2.

### Yeast culture

Unless otherwise stated, all strains were grown overnight in Yeast extract Peptone Dextrose (YPD) media at 30°C with shaking, washed 3x with H_2_O, diluted to an OD_600_=0.1 in SC-Met (synthetic complete media without methionine), further grown for 2.5 h at 30°C and assayed as required in early logarithmic phase. Plasmid-containing strains were grown overnight in SC-Ura (synthetic complete media without uracil), washed 3x in H_2_O and grown for 2.5 h in SC-Ura-Met media. Unless otherwise indicated, Mup1 endocytosis was triggered by addition of 1 mM methionine to the media. SC all: synthetic complete media. All media were supplemented with 2% glucose.

### Microscopy and imaging

Epifluorescence, Fluorescence Recovery After Photobleaching (FRAP) and Total Internal Reflection Fluorescence Microscopy (TIRFM) were performed on an fully automated iMIC stand (FEI/Till Photonics) with an Olympus 100X 1.45 NA objective. DPSS lasers (75 mW) at 491 nm (Coherent Sapphire) and 561 nm (Cobolt Jive) were selected through an acousto-optical tunable filter. A two-axis scan head was used to adjust TIRFM incidence angles or FRAP positions. An additional galvanometer was used to switch between illumination paths. Images were collected with an Andor iXON DU-897 EMCCD camera controlled by the Live Acquisition (Till Photonics) software. For two-color TIRFM, incidence angles were adjusted separately for each laser. Separate filters were used for detection of green and red fluorophores. 4-color 100 nm microspheres (ThermoFisher Sci.) were used to determine the Point Spread Function (PSF) in both channels and to correct for the offset between filters. For two-color TIRFM, coverslips (Knittel Glass No. 1) were cleaned by sonication in absolute ethanol (Sigma), > 99.5% acetone (Sigma), 1 M NaOH (Roth), ddH_2_O and finally stored in absolute ethanol. Long term imaging experiments were performed using glass bottom culture dishes (MatTek). To immobilize cells, coverslips and glass bottom dishes were coated with 1 mg/ml concanavalin A (Sigma).

### Image processing and analyses

Images were processed using Fiji and Matlab (Mathworks Inc., Natick, MA). Images were contrast-adjusted and zoomed for presentation purposes in Figures only. For image cleanup and denoising we routinely used the background subtraction algorithm in Fiji (radius 50 pixel). Kymographs and intensity plots were created using the corresponding features in Fiji. All TIRFM images were deconvolved using the Lucy-Richardson algorithm in MATLAB (deconvlucy), with PSF functions calculated from 4-colour 100 nm microsphere images and 12 iterations. Protein colocalization (Pearson Mean), GFP intensity distribution (Network factor) and GFP overlap were calculated from TIRFM images using a customized automated script and gui written in MATLAB. Cells were automatically detected from blurred raw images, deconvolved and thresholded using an adaptive filter based on the difference between raw and deconvolved images. Masks were then generated for each separate channel. Calculations were based on combined (AND) masks (Figure S1). Additional filters based on object size and intensity were used to discard false detections. Colocalization between GFP and RFP signals was determined by calculating the linear correlation coefficient (given as Pearson Mean) of the fluorescence within the generated masks. The GFP overlap indicates the fraction of GFP signal within the RFP mask and was used to better characterize co-localization of highly clustered proteins. The Network factor represents the variance of the intensity distribution and was determined for entire cells (no masks) detected in deconvolved images. A low network factor represents highly clustered structures (patches) while higher values correspond to network-like distributions. All images shown were scaled 3x and color-coded using Fiji. TIRFM and equatorial images in Figure 7A-C were processed using the Super-Resolution Radial Fluctuations (SRRF) plugin in Fiji^41^.

### PM expression and endocytosis measurements

All measurements are given as ratio between fluorescence intensity at the PM and the cytosol. Intensity values were determined from automatically-detected cells using a customized MATLAB script. Cells expressing no PM fluorescence were quantified manually in Fiji. The PM intensity distribution in Figure 1B was calculated manually for strains taken from the UCSF GFP collection^37^ (Table S2). The endocytosis assay in Figure 1D was performed at 22°C and the values automatically-determined on completely static cells throughout the experiment using the detected PM and cytosol areas at t = 0 min.

### Ubiquitination assay and Western blot

Strains were initially grown in SC-Met as described under “Yeast Culture”. Aliquots of 1 ml cells (OD_600_=1) were then supplemented with 1 mM Met for the indicated time. Cells were centrifuged at 16.000 g for 30 s, the supernatants removed and the pellets quickly transferred to liquid nitrogen to stop ubiquitination. Cell pellets were then suspended in 100 μl of ice-cold protease-inhibitors (2 mM PMSF, 2x complete protease inhibitors, 8 mM EDTA). Next, 50 μl of a 2 M NaOH solution were added, cells incubated for 10 min on ice, 50 μl of a 50% TCA-solution added and cells again incubated for 10 min on ice. Finally, samples were centrifuged for 5 min at 18.000 g (4°C) and the pellets suspended in 50 μl sample buffer 1 (100 mM Tris-HCl pH 6.8; 4 mM EDTA; 4% SDS; 20% Glycerol; 0.02% Bromophenol blue and 50 μl sample buffer 2 (1 M Tris; 2% β-mercaptoethanol. The temperature was kept at 4°C throughout the entire protein extraction. Samples were loaded and separated using standard SDS (8.17% polyacrylamide / 13.3% glycerol) gel electrophoresis. Proteins were transferred to a nitrocellulose membrane and probed with monoclonal mouse IgG1κ anti-GFP (Roche, #11814460001) and monoclonal anti-GAPDH (abcam, #ab125247). Primary antibodies were detected with peroxidase AffiniPure Goat Anti-Mouse IgG secondary antibodies (Jackson ImmunoResearch Inc., #115-035-003) followed by chemiluminescence detection with Lumi-Light^Plus^ Western Blotting Substrate (Roche). Signals were detected with an advanced fluorescence imager (Intas).

### Uptake measurements

Mup1 transport activity was analyzed by measuring the uptake of ^14^C-methionine (specific activity: 1417484 nmoles/cpm, final concentration 40 μM, Perkin-Elmer, Boston, MA) 20, 40, 60 and 80 seconds after start of the reaction. Accumulated counts were measured using a β-counter (LS 6500, Beckman Coulter Inc., Brea, CA) and line-fitted to determine the uptake rate (given in nmol/min per mg of total protein). All strains were grown in SC-Met as described under “Yeast culture” prior to the uptake measurement. All R^2^ > 0.95. Further details as described in ^42^.

### Growth assays

Yeast growth assays were performed on plates in the presence of the toxic methionine analog selenomethionine (SeMet, Sigma-Aldrich) or the toxic arginine-related non-proteinogenic amino acid Canavanine (Sigma-Aldrich). Plates were prepared as required in SC all, SC-Met or SC-Arg and the corresponding drug. Cells were grown in SC-Met as described under “Yeast culture” with the exception of final 2.5 h growth on SC-Arg for the canavanine-based assay. Cells were spotted on plates in a 5x dilution series, starting at an OD_600_ = 0.1. Cells were then grown at 30°C for 3 days (SC all and SC-Arg) and 5 days (SC-Met). Drug concentrations used as indicated in the respective figures.

### Sequence alignments and tertiary structure prediction

Sequence alignments were performed using the open source Clustal Omega (EMBL-EBI) software and color-coded as indicated in the respective Figure. Mup1 tertiary structure was predicted by the Protein Homology/analogY Recognition Engine V 2.0 (Phyre2)^43^. The C-terminal “plug” was inferred based on the predicted homology towards the bacterial transporter GadC^31^. Final structures were color-coded using the Java-based free Protein Workshop software^44^.

### Inhibitor experiments

Myriocin (Myr, Sigma-Aldrich) and aureobasidinA (AbA, Clontech) were used to inhibit sphingolipid synthesis. Myr inhibits the serine palmitoyltransferase (SPT), which catalyzes the first step in the sphingolipid biosynthetic pathway. AbA inhibits the yeast inositol phosphorylceramide synthase, which catalyzes a late step in the biosynthesis of complex sphingolipids. Both drugs were dissolved in 100 % ethanol at 5 mM and stored at 4°C, used at a final concentration of 5 μM and incorporated into the cell suspension for 1 h at 30°C with shaking before imaging. The macrolide compound rapamycin (Rap, Santa Cruz) was used to inhibit TORC signaling, dissolved in DMSO, used at a final concentration of 1 μg/ml and incorporated into the cell suspension for 30 min at 30°C with shaking before imaging. The *tor1-2* mutant^40^ makes the TORC1 complex rapamycin-insensitive while the *avo3*ΔC mutant renders the TORC2 complex rapamycin-sensitive^25^. Latrunculin B (LatB, Enzo Life Sciences) was used at 100 μM to inhibit actin assembly in endocytosis experiments.

### 3D Structured illumination microscopy (3D-SIM)

3D-SIM was performed on a Nikon Ti-E N-SIM / N-STORM setup. The light source was controlled by a LU-NV Laser Unit with a 488 nm (GFP) and a 561 nm (RFP) laser line using independent N-SIM filter cubes. The objective was a CFI SR APO TIRF 100xH oil, NA 1.49, WD 0.12 mm (Nikon). Patterns were generated with an optimized grating (3D SIM 100x, 1.49 405-640 nm (Ex V-R)) and the images collected with an Andor Ultra EM-CCD and DU-897 Camera controlled by the NIS-Elements (Nikon) software. 4-color 100 nm microspheres (ThermoFisher Sci.) were used to determine the PSF using the NIS-Elements software. Z-stacks were taken as 0.2 μm steps from the cell surface close to the glass coverslip up to 1 μm into the cell. Image z-Stack-Reconstruction was conducted automatically with default settings by the NIS-Elements software. The samples were prepared as for TIRFM imaging. The coverslips were from VWR (No. 1.5) and the immersion oil from Nikon (nd: 1.515). Images represent maximum projections of 9-11 planes.

### GFP trap system

In order to alter the localization of Mup1 we endogenously fused the GFP binder (GB) to the C-terminus of the protein of interest. The GB is a single-chain, high-affinity GFP antibody from camelids^45^ that binds its target when both GFP and GB are co-expressed.

### Statistics

Mean values, standard deviations (SD), standard error of the mean (SEM) and numbers of measurements (n) are provided for all quantified results. All quantifications are summarized in Table S1. Values were always pooled from at least three independent experiments except for Met uptake experiments (more than 2). All replicates are biological replicates. Error bars in graphs are explained in the respective legends. Graphs were prepared in Prism (GraphPad). Significance values in Figure 6E were determined using the non-parametric Wilcoxon-Mann-Whitney test. p-values are given in the figure legend.

## Acknowledgements

We thank P. Hardy, S. Wedlich and members of the Wedlich-Söldner lab for valuable comments on the manuscript. We are grateful to C. Gournas and B. André for help with Met uptake assays. This work was supported by the German Research Foundation (SFB944, 1348, WE2750/4-1, RWS) and the Cells-in-Motion Cluster of Excellence (EXC1003–CiM, University of Münster, RWS). JVB was supported by a Postdoctoral Fellowship from the Basque Government. The authors declare no competing financial interests.

## Author Contributions

JVB, FS and RWS conceived the project, designed and conducted experiments, and analyzed data. JVB, FS, AE and DH conducted experiments and analyzed data. JK and MSH wrote Matlab code and analyzed data. JVB and RWS wrote the manuscript with the help of all authors.

---

**Figure S1.**
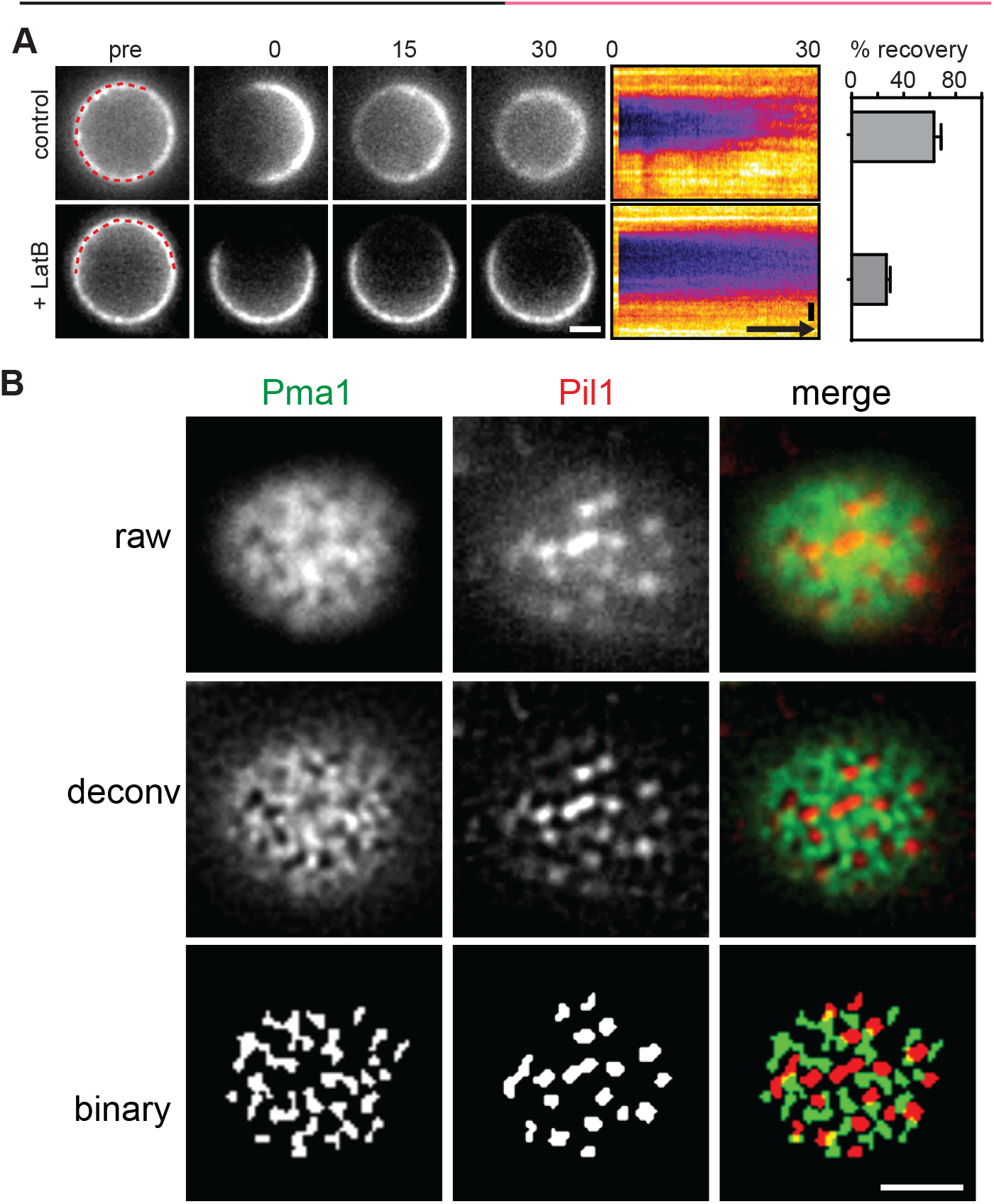
(**A**) FRAP analysis of Mup1-GFP in the presence and absence of 100 μM LatB. Times are given in min relative to localized bleaching of fluorescence. The kymograph was drawn around the indicated cell periphery (red dashed lines) and shows fluorescence recovery in the bleached area. Mup1-GFP fluorescence recovery was determined in the bleached area at t = 30min. Time arrow represents 10 min. Values are means ± SD, n = 10 cells. (**B**) Representative raw, deconvolved (deconv) and thresholded (binary) TIRFM images generated by an automated Matlab algorithm to determine the degree of colocalization of GFP and RFP signals (Pearson Mean) and to quantify the fraction of the GFP signal present in the RFP-labelled structure (GFP overlap). Scale bar: 2 μm. Values are listed in Table S1.

**Figure S2.**
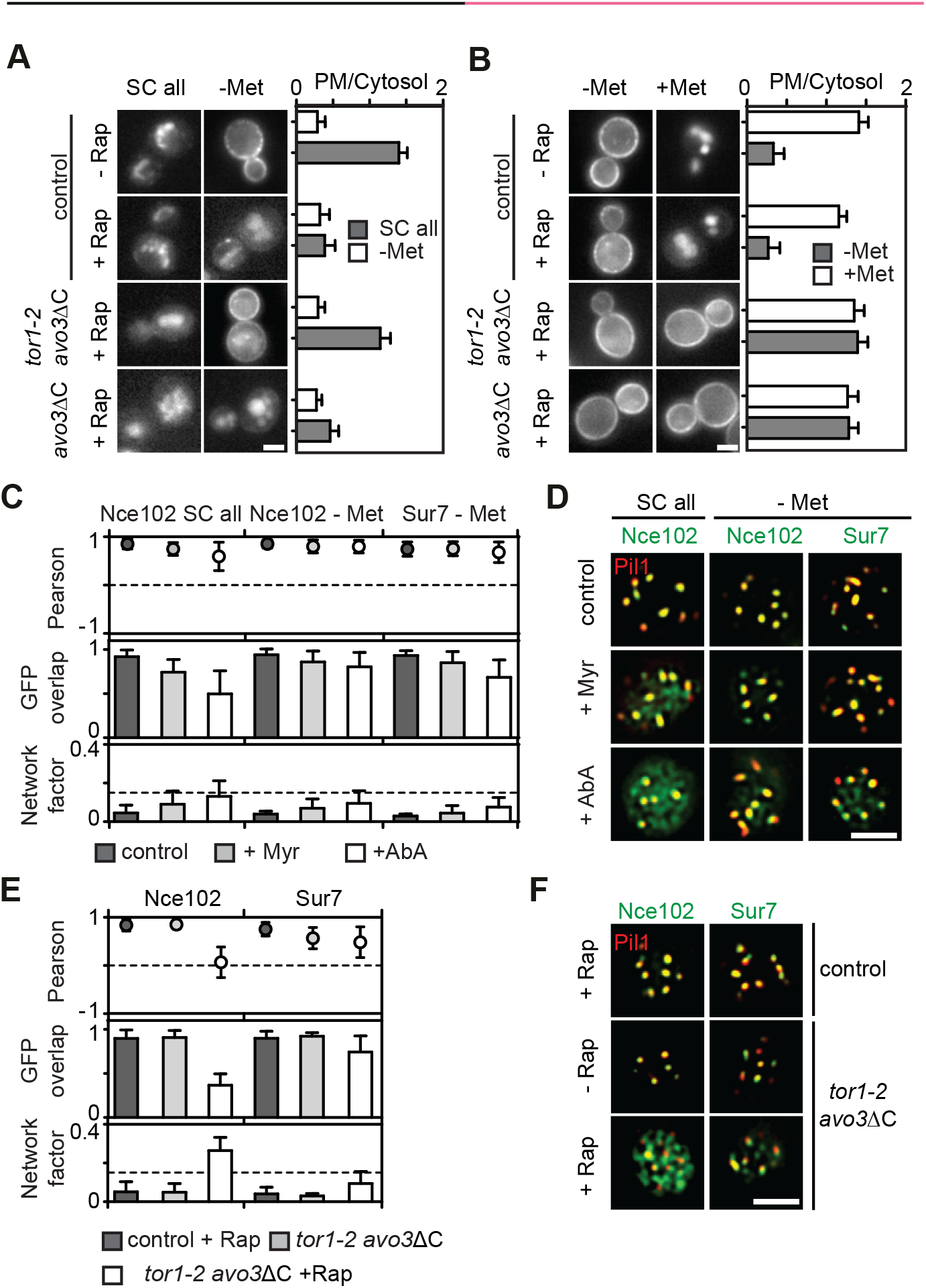
(**A, B**) Requirements of TORC1 and TORC2 for the delivery of Mup1-GFP to the PM upon methionine starvation (**A**) and for Mup1 endocytosis upon addition of methionine (**B**). In control cells only TORC1 is rapamycin (Rap) sensitive, in the *avo3*ΔC mutant both TORC1 and TORC2 are sensitive to Rap, and in the Tor1-2/*avo3*ΔC mutant only TORC2 is Rap-sensitive. Representative equatorial images are shown. (**C**) Influence of sphingolipid stress on the lateral segregation of the tetraspanners Nce102 and Sur7 under Met starvation. Pil1-RFP was used to determine the degree of concentration within the MCC. (**D**) Representative two-color TIRFM images from the experiments summarized in C. (**E**) Nce102 leaves the MCC upon TORC2 inhibition. (**F**) Representative two-color TIRFM images from the experiments summarized in E. All values plotted are means ± SD, n = 20-150 cells. Scale bars: 2 μm. All measured values are listed in Table S1.

**Figure S3.**
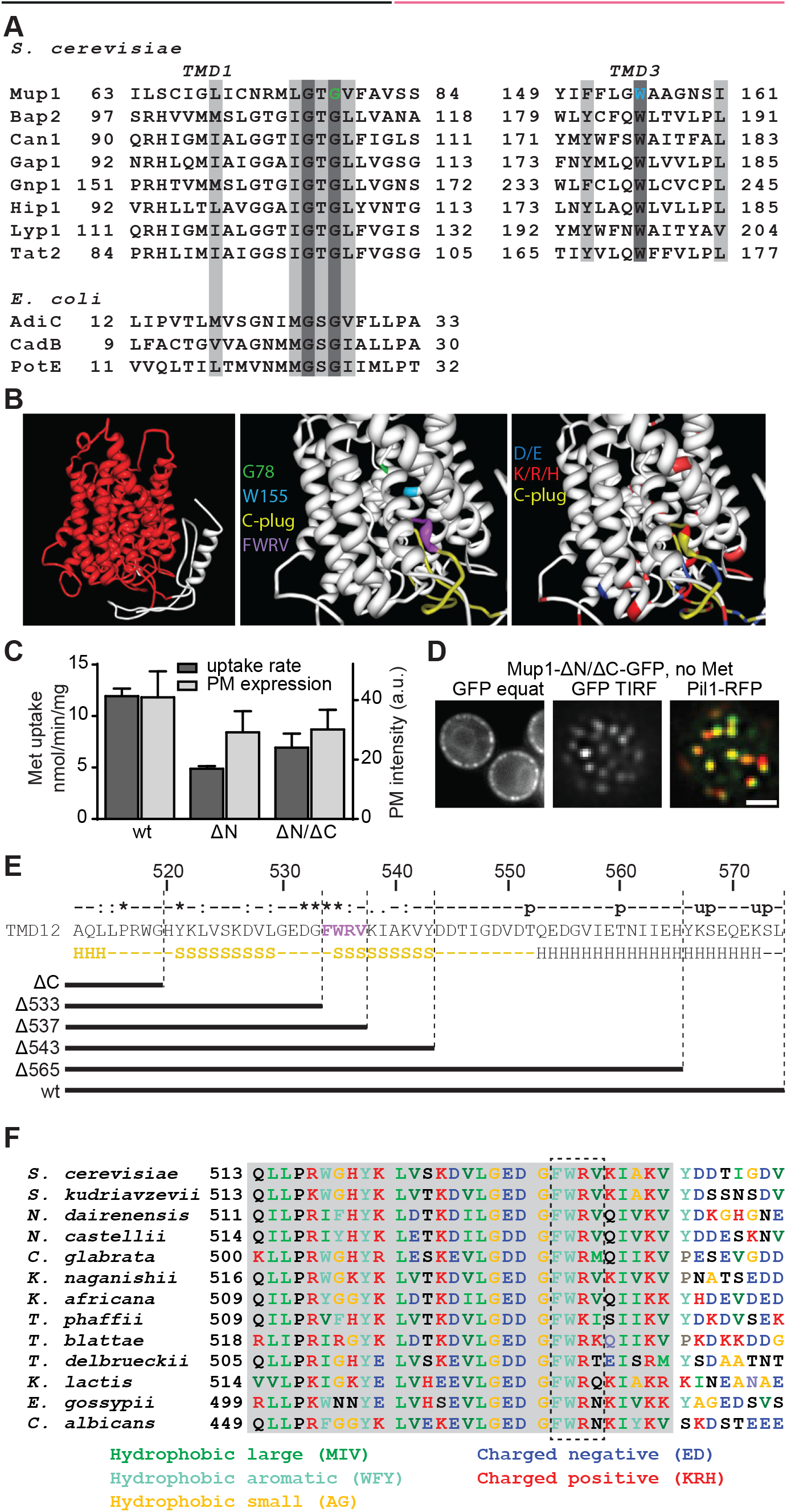
(**A**) Sequence alignments of predicted TMD1 and TMD3 of Mup1 with those of other yeast and *E. coli* transporters. Conserved residues (dark grey), conservative replacements (light grey) and selected Mup1 mutations (G78 and W155) are highlighted. (**B**) Structural organization of Mup1 as predicted by the Phyre2 software^43^. High (red) and low (white) confidence predictions are highlighted (left panel). Mutations affecting Met transport (green and light blue), the C-terminal “plug” oriented towards the substrate binding pocket (yellow), the conserved C-terminal motif required for Mup1 localization and function (purple), as well as negatively (dark blue) and positively (red) charged residues around the binding pocket and the “C-plug” are indicated (middle and right panels). (**C**) ^14^C-methionine uptake rates and PM expression of terminally-truncated and GFP-fused Mup1 mutants. Values are means ± SD, n = 2 experiments (uptake) and n > 70 cells (PM expression). (**D**) Representative equatorial and TIRFM images of Mup1ΔN/ΔC in the absence of Met, showing its delivery to the PM and partitioning into the MCC (colocalization with Pil1-RFP). (**E**) Sequence of the Mup1 C-terminal region. Degrees of conservation among Mup1 homologs (from ClustalW) calculated from various other fungi as shown in F (u: predicted ubiquitination site, p: predicted phosphorylation site) are indicated above the sequence, with secondary structure predictions (H: alpha helix, S: beta sheet) below it. The predicted “C-plug” is indicated in yellow. The different C-terminal truncations analyzed are shown. (**F**) Sequence alignment of the Mup1 C-terminal segment (513-550) with homologs from various other fungi. Scale bar in D: 2 μm.

**Figure S4.**
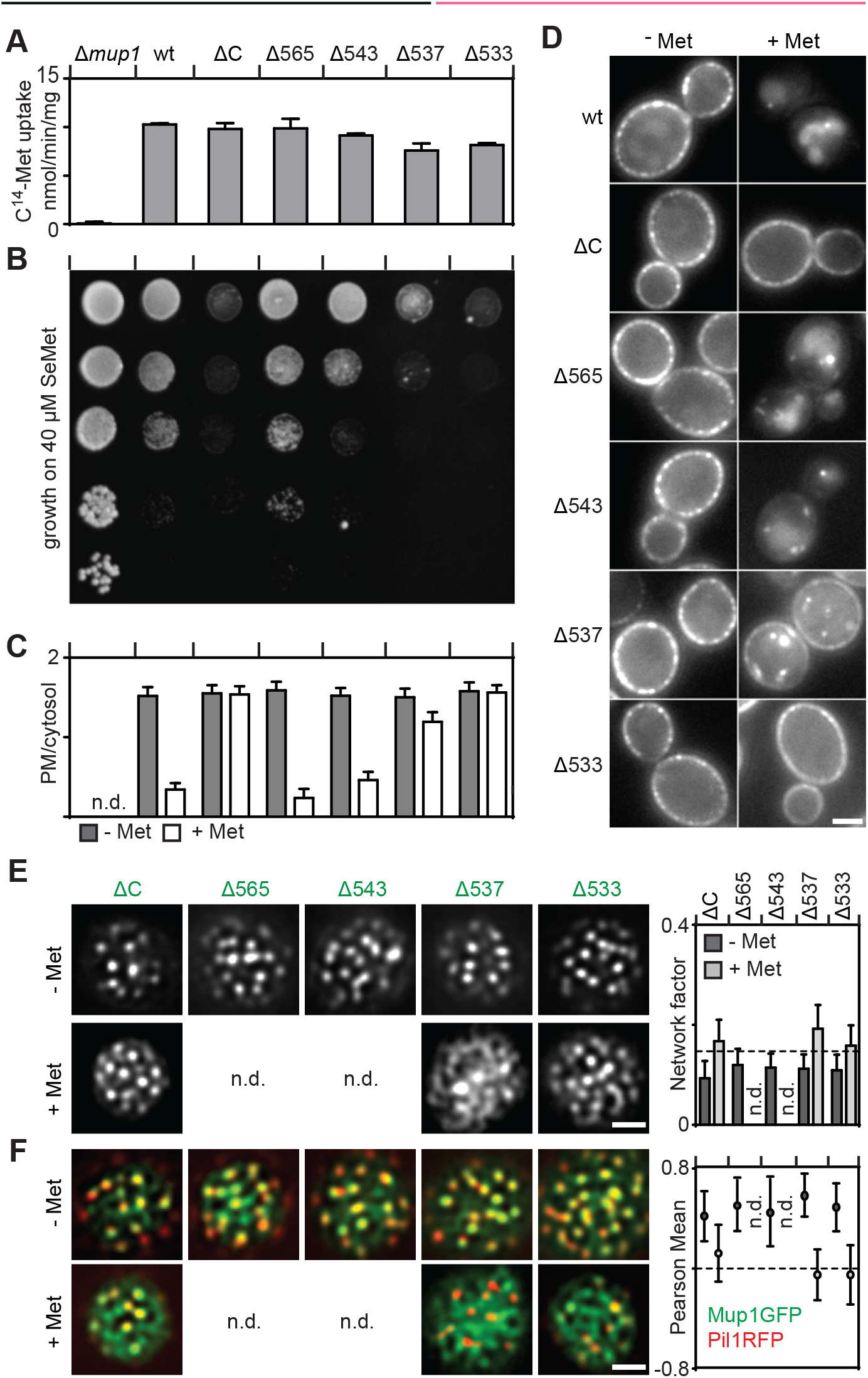
Effects of various C-terminal truncations on the function, lateral PM segregation and turnover of Mup1. (**A, B**) Function of different Mup1 mutants as measured by direct quantification of ^14^C-Met uptake (**A**) and growth sensitivity to the toxic Met analog selenomethionine (SeMet, **B**). (**C, D**) Endocytic internalization of different Mup1 mutants. Ratios of PM to cytosolic fluorescence intensities (**C**) and representative images at equatorial planes (**D**) are shown. (**E, F**) Lateral PM segregation shown in representative TIRFM images and quantification of the network factor (**F**) or the colocalization with Pil1-RFP (**G**). Mutants utilized: wt (wild-type Mup1), ΔC (deletion of C-terminus after aa519), W155A and G78N (respective point mutants), Δ565/543/537/533 (deletion of C-terminus beyond the indicated position). All strains refer to Mup1 variants fused to C-terminal GFP. Values are means ± SD, n = 2-4 experiments (A) and n = 50-200 cells (C, E, F). n.d.: not determined. Scale bars: 2 μm. All values are listed in Table S1.

**Figure S5.**
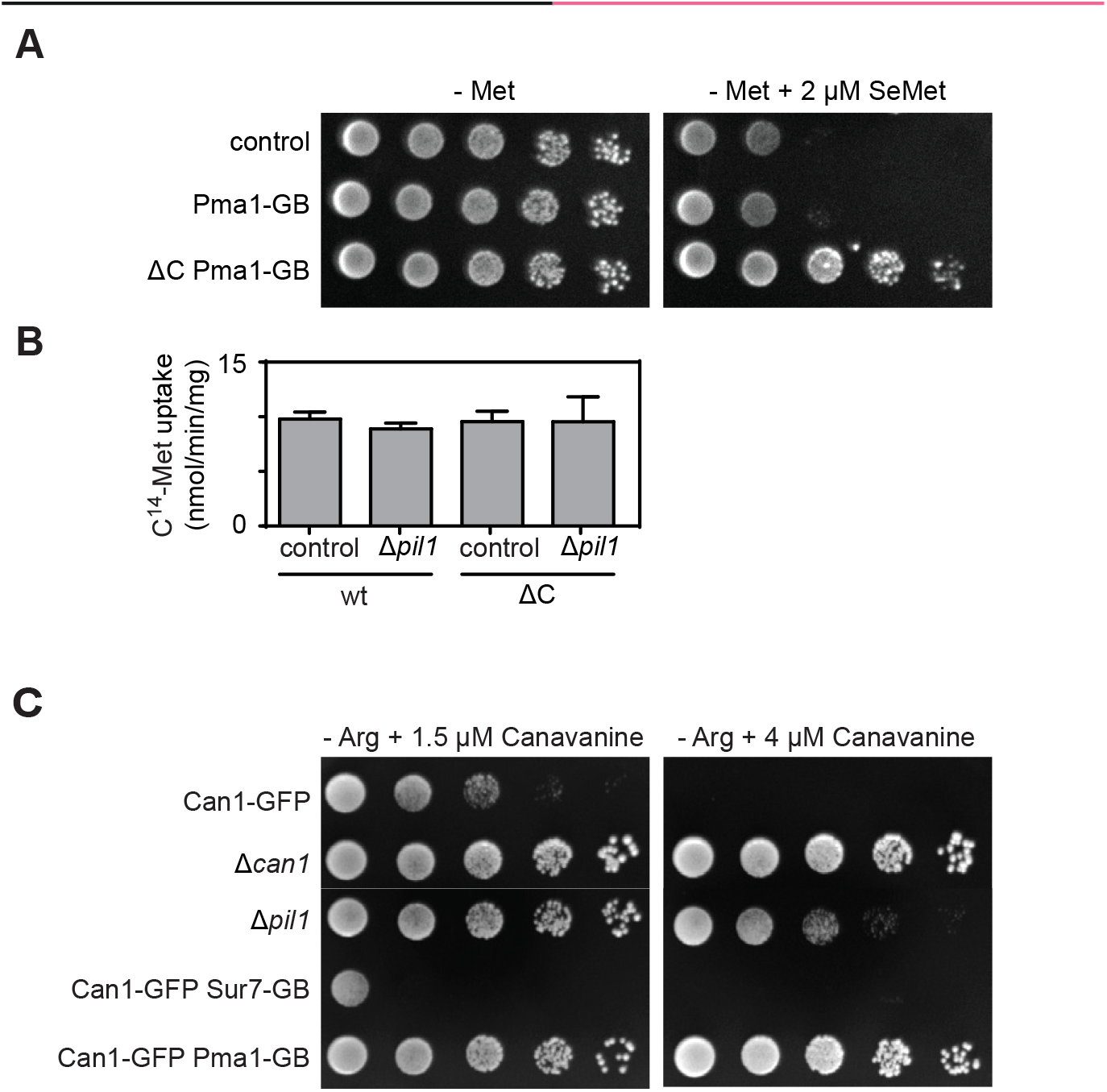
(**A**) Growth assay of indicated mutants in the absence of methionine and in the absence or presence of 2 μm SeMet. (**B**) ^14^C-methionine uptake rate measured for Mup1 and ΔC in the wt and the Δ*pil1* background. Values are means ± SD, n = 2-4 experiments. (**C**) Growth assay of indicated mutants in the absence of arginine and the presence of indicated concentrations of the toxic arginine analog canavanine. All values are listed in Table S1.

**Movie S1.** Actin patches assemble on Mup1 clusters. TIRFM movie showing assembly of endocytic patches marked by the actin binding protein Abp1-RFP at the PM of yeast cells expressing Mup1-GFP. The movie starts 10 min after addition of Met to induce MCC exit and ubiquitination of Mup1. Examples of actin patch assembly that remove Mup1 signal from the PM are marked with asterisks. Channels were separately filtered using the SRRF plugin in Fiji (see methods). Scale bar: 2 μm. Frames were taken every 5 s.

